# Conformational Tuning Shapes the Balance between Functional Promiscuity and Specialization in Paralogous *Plasmodium* Acyl-CoA Binding Proteins

**DOI:** 10.1101/2023.05.12.540612

**Authors:** Rahul Dani, Westley Pawloski, Dhruv Kumar Chaurasiya, Nonavinakere Seetharam Srilatha, Sonal Agarwal, David Fushman, Athi N. Naganathan

## Abstract

Paralogous proteins confer enhanced fitness to organisms *via* complex sequence-conformation codes that shape functional divergence, specialization, or promiscuity. Here, we resolve the underlying mechanism of promiscuous binding *versus* partial sub-functionalization in paralogs by studying structurally-identical Acyl-CoA Binding Proteins (ACBPs) from *Plasmodium falciparum* that serve as promising drug targets due to their high expression during the protozoan proliferative phase. Combining spectroscopic measurements, solution NMR, SPR and simulations on two of the paralogs, A16 and A749, we show that minor sequence differences shape nearly every local and global conformational feature. A749 displays a broader and heterogeneous native ensemble, weaker thermodynamic coupling and cooperativity, enhanced fluctuations, and a larger binding-pocket volume, compared to A16. Site-specific tryptophan probes signal a graded reduction in the sampling of substates in the *holo* form, which is particularly more apparent in A749, hinting at conformational-selection-like mechanism of binding. The paralogs exhibit a spectrum of binding affinities to different acyl-CoAs with A749, the more promiscuous and hence the likely ancestor, binding 1000-fold stronger to Lauroyl-CoA under physiological conditions. We thus demonstrate how minor sequence changes modulate the extent of long-range interactions and dynamics, effectively contributing to the molecular evolution of contrasting functional repertoires in paralogs.

## Introduction

Gene duplication events are key drivers of evolution wherein a gene is copied within an organism’s genome, resulting in two copies of the gene. This can happen through various mechanisms, such as retro-transposition, in which the gene is transcribed into RNA and then reinserted into the genome, or homologous recombination, in which the gene is copied through exchange with a similar gene on a homologous chromosome. The copies of a duplicated gene, known as paralogs, often retain the same function as the original gene and contribute to organismal fitness *via* dosage selection, but may also undergo functional specialization, diversification, and loss of activity followed by re-functionalization.^1–4^ It is possible that one copy of the gene maintains its original (ancestral) function while mutations accumulate in another copy contributing to neo-functionalization or sub-functionalization. Neo-functionalization occurs when one of the duplicated genes acquires a completely new function, often through the acquisition of new regulatory elements or the accumulation of mutations that alter the protein’s function. Sub-functionalization occurs when the duplicated genes acquire partially overlapping functions, leading to a division of labor between the two copies.^5–12^

Gene duplication and the evolution of paralogous proteins therefore provide a mechanism for organisms to adapt to changing environments and to develop new traits. However, the maintenance of duplicated genes and the evolution of paralogous proteins also come with costs, such as the increased demands on energy and resources. Paralogous proteins often have similar sequences and structures, with the differences arising due to the requirement of functional specialization-cum-diversification. Although both these features are expected, the mechanism through which such functional advantages are realized by subtle changes in the sequences are less understood. This is primarily due to the nature of the intra-protein interaction network that can be wired in different ways (*i.e.*, the interaction network can be tuned to similar extents *via* different mutations) to achieve the same or different functional outputs through modulation of dynamics in the native ensemble.^13–18^ Although sequence alignments could highlight the degree of conservation, it does not provide insights on the conformational aspects that are perturbed and their extents, which effectively determine the functional output.

Here, we investigate the molecular determinants of partial sub-functionalization in paralogous proteins by studying acyl-CoA binding proteins (ACBPs) from the protozoan *Plasmodium falciparum*. ACBP is an α-helical cytosolic protein which is highly conserved among all eukaryotic species ranging from protists to mammals,^19, 20^ and a popular model system to study folding mechanisms.^21–27^ It binds small-to long-chain fatty-acyl Coenzyme A (CoA), protects them from hydrolases, and transports them to mitochondria for β-oxidation, thus serving as a crucial player in lipid metabolism.^28^ In addition, ACBPs donate acyl-CoAs for synthesizing variety of lipid derivatives like triacyl glycerol, ceramide and ultra-long chain fatty acids.^29^ ACBP is also found in the nucleus where it binds to the lipid binding domain of HNF-4α with high affinity and alters its conformation such that it can recruit SRC1/p300 coactivators and regulate expression of genes related to lipid metabolism.^30^

*P. falciparum* harbors three isoforms of ACBPs – A16, A749 and A99 – that show high expression during the trophozoite and schizont stages and hence play a critical role in the replication cycle of the parasite.^31, 32^ The presence of three ACBP paralogs highlights an evolutionary selection for function, which is not fully understood. While only the structure of the *holo* A749 is available,^33^ AlphaFold2 predictions^34, 35^ reveal that the 3D structures of the paralogs are near-identical with an overall mean C_α_-RMSD of 0.3 Å (Figure 1A). This is not surprising as the sequences themselves are highly similar (75%; Figure 1B). The *P. falciparum* ACBPs (pfACBPs) exhibit the classic four-helix bundle topology, with the exposed surfaces of the three helices H1, H2 and H3 forming a binding pocket to accommodate acyl-CoA ligands. A closer look at the sequences points to differences concentrated in H1, the loop connecting H1 and H2 (L1), and the apparently unstructured C-terminal tail. Also, the C-terminal tail harbors a conserved tryptophan 86 (W86), whilst in other members it is predominantly tyrosine that occupies this position (Figure S1).

**Figure 1.**
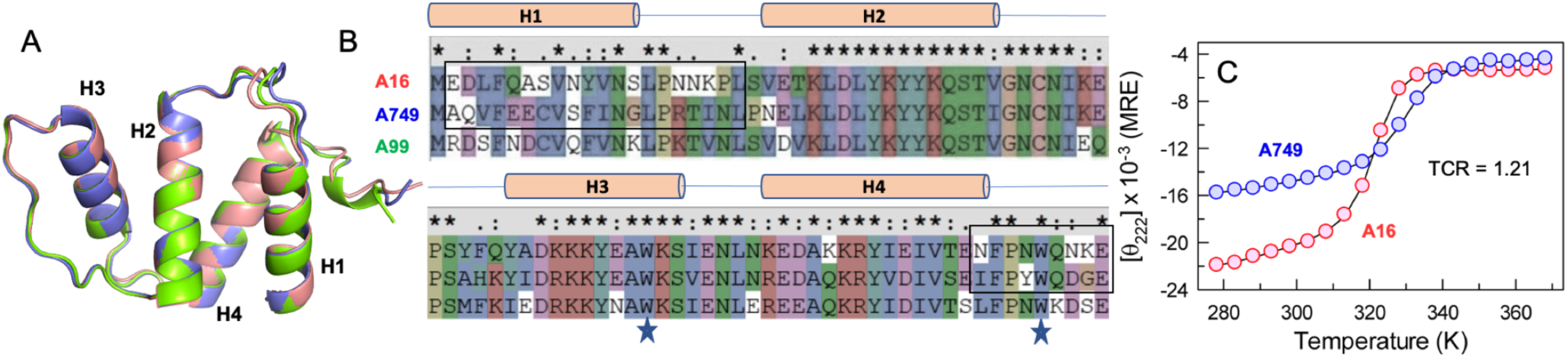
Structural and sequence comparison. (A) Superimposed 3D structures of *Plasmodium falciparum* ACBP paralogs. (B) Sequence alignment from ClustalX2^36^ with the boxes highlighting the positions that display maximal differences. Stars indicate the position of tryptophan residues W60 and W86. (C) Thermal unfolding curves monitored by far-UV CD at 222 nm reported in mean residue ellipticity units of deg. cm^2^ dmol^−1^. The black curve through the data points is a two-state fit. TCR is the thermodynamic cooperativity ratio (see text for details).

Given these observations as the starting point, we perform an array of spectroscopic experiments on two of the members of the pfACBPs, A16 and A749, both in the *apo* and *holo* forms to extract and understand the extent of differences in stability, cooperativity, and local-global dynamics. We validate our experimental observations *via* statistical modeling and MD simulations, followed by functional studies using a collection of acyl-CoA ligands of different carbon chain lengths to explore the extent of functional specialization. Finally, we discuss the implications of our work for understanding the sequence-dynamics-function relation in proteins and the role of long-range interactions in mediating partial sub-functionalization in paralogs.

## Methods

### Purification of ACBP paralogs and mutants

Freshly transformed BL21(DE3) cells containing the gene sequence corresponding to the proteins A16 (MKEKEMEDLFQASVNYVNSLPNNKPLSVETKLDLYKYYKQSTVGNCNIKEPSYFQYADKKKYEAWKSIENLNKEDAKKRYIEIVTENFPNWQNKEEKEK) or A749 (MKEKEMAQVFEECVSFINGLPRTINLPNELKLDLYKYYKQSTIGNCNIKEPSAHKYIDRKKYEAWKSVENLNREDAQKRYVDIVSEIFPYWQDGEEKEK) in the pTXB1 vector were grown in 2 L LB media containing 100 μg/mL ampicillin. The culture was induced with 1 mM IPTG at an OD_600_ of ~1.0. Following induction, the culture was incubated for 4 hours at 37 °C, cells were harvested by centrifugation, and the pellets were stored at −80 °C. The cell pellet corresponding to ~1.3 L culture broth was dissolved in 80 mL of lysis buffer (20 mM Tris, 500 mM NaCl, 1 mM EDTA, pH 8.5) containing 1 mM PMSF, lysed, and centrifuged at 10,500 RPM for one hour at 4°C. The resulting supernatant was loaded onto a chitin resin column (New England Biolabs Inc.) previously equilibrated with same buffer at the flow rate of 0.5 mL/min. Following this, the column was washed with the same buffer for 10 column volumes (CV). Finally, buffer containing 50 mM β-mercaptoethanol was passed through the column for initiation of the cleavage reaction. Following a 36 hour incubation at room temperature, the protein was eluted using the same buffer. The eluted protein was passed through a size exclusion chromatography (SEC) column (HiLoad 26/600 Superdex 75pg, Cytiva) previously equilibrated with 150 mM ammonium acetate buffer at pH 8.0. Eluted fractions containing purified protein (as checked using SDS-PAGE) were pooled and lyophilized. W60A and W86A mutants of A16 and A749 were purified using the same protocol as above. For generating ^15^N labeled proteins, a single colony of freshly transformed BL21(DE3) cells with pTXB1 plasmid containing wild-type protein or mutant gene sequence was inoculated in 1 L of M9 minimal media containing one gram of ^15^NH_4_Cl as the sole source of nitrogen. The culture was grown at 37°C and 180 RPM till the OD at 600 nm reached ~1.0. Following this, culture was induced with 1 mM IPTG and grown at 37°C for 4 more hours after which the cells were harvested by centrifugation and the pellet was stored at −80°C until sonication. Further purification procedure is same as unlabeled proteins as mentioned above.

### Spectroscopy and stopped-flow kinetics

Far- and near-UV CD spectra were acquired at protein concentrations of ~18 μM and ~70 μM, respectively, in degassed 40 mM sodium acetate buffer, pH 4.0 (ionic strength adjusted to 100 mM using NaCl). Spectra were recorded on a Jasco J-815 spectrophotometer coupled to a Peltier system. The resulting signal was converted to mean residue ellipticity (MRE) units of deg. cm^2^ dmol^−1^. Fluorescence emission spectra on excitation at 295 nm wavelength were recorded in a Chirascan^TM^-plus qCD instrument (Applied Photophysics) coupled to a Peltier system (Quantum Northwest Inc.) at a sample concentration of ~8-10 μM. Excitation and emission monochromator bandwidths were set to 5 nm. Time-resolved fluorescence measurements were performed as detailed before in a ChronosBH (ISS Inc.) spectrometer on excitation with a 300 nm LED.^37^ All decay curves were recorded until the peak count reached 10^4^ or the total count approached 10^8^. The traces were best reproduced by tri-exponential functions with the chi-square values being <1.2 at all temperatures.

Unfolding and refolding traces were recorded at 285K using a Chirascan SF3 Stopped-flow setup (Applied Photophysics) with dead time of ~2-3 millseconds. The protein sample was excited with 280 nm LED source and emission signal was collected through a 295 nm filter. 16 kinetic traces were taken for every sample at one-minute intervals, averaged, and fit to single- or bi-exponential functions. For the refolding kinetics, proteins were dissolved in 6.5 M urea and kinetic traces were recorded at final urea concentrations of 0.6 to 4.2 M.

### Differential scanning calorimetry

Apparent heat capacities were measured in a Microcal VP-DSC (Malvern, UK) at protein concentrations ranging from ~60 μM to ~125 μM, and at a scan rate of 1.5 K/min. Lyophilized ACBPs were dissolved in 100 mM ionic strength, pH 4 buffer and buffer-exchanged employing a 26/10 HiPrep Sephadex G desalting column. Several buffer-buffer heat capacity scans were acquired before and after the protein scans to ensure proper equilibration of the cells. Samples were equilibrated for 20 minutes at 278 K before acquisition of scans. Absolute heat capacity was derived from the concentration dependence of apparent heat capacity as outlined by Sanchez-Ruiz and co-workers.^38^ For the experiments in the presence of the ligand, a stock solution of Stearoyl-CoA (C18-CoA) was prepared in the experimental buffer. Following this, the ligands were added to protein solutions to obtain a protein:ligand molar ratio of 1:0.2. The heat capacity of proteins in the presence of ligand were corrected for any potential ligand effects by subtracting them with the specific heat capacity values of the protein-free ligand solutions.

### Nuclear magnetic resonance

The NMR measurements were performed for a range of temperatures, from 286 K to 330 K, on Bruker Avance III 600 MHz and 800 MHz spectrometers equipped with cryogenic probes. The NMR sample contained ^15^N-enriched A16 or A749 WT, and W86A/W60A variants of A749, dissolved in a 40 mM sodium acetate buffer (pH 4.0) containing 60 mM NaCl, 10% D_2_O, and 0.02% (v/v) NaN_3_. A freshly prepared sample was used for each set of measurements. Heteronuclear experiments included ^1^H-^15^N HSQC, SOFAST-HMQC,^39^ CEST,^40^ and CPMG-based 15N transverse relaxation dispersion measurements,^40^ which were typically recorded with the spectral widths of 9615-12820 Hz (^1^H) and 2128-2920 Hz (^15^N). All spectra were processed in NMRPipe^41^ and analyzed with CCPNMR V2.^42^ The ^1^H and ^15^N chemical shifts were adjusted to the temperature dependent lock shift during processing.

### Surface plasmon resonance (SPR)

The ligand binding experiments were carried out using a Biacore 3000 instrument. pfACBPs were immobilized onto CM5 sensor chips using an amine coupling kit. The channels were equilibrated with either pH 4 buffer (100 mM ionic strength) or PBS (phosphate buffered saline), pH 7.4 at 298 K. The first channel on the chip was kept blank to monitor and subtract for any potential artifacts from buffer or analyte injections. Various concentrations of the analyte - Hexanoyl-CoA (C6), Lauroyl-CoA (C12) or Stearoyl-CoA (C18) - were prepared in the same buffer. 50 µL of analyte solutions were injected at a flow rate of 30 µL/min followed by monitoring the dissociation phase for at least 300 seconds. Buffer injections were routinely given after every analyte to ensure baseline stability. In some cases when the analyte is tightly bound to the protein (especially for the longer carbon-chain acyl-CoAs), the channel was regenerated using 2 M MgCl_2_. The sensograms for a particular protein and for a specific ligand identity (C6-, C12- or C18-CoA) at different ligand concentrations were globally fit to a 1:1 binding model using BIAevaluation 3.0 software to estimate the on-rate for association (*k_on_*), the off-rate of dissociation (*k_off_*), and hence the dissociation constant (*K_D_*).

### Statistical mechanical modeling

An in-depth description of the Ising-like Wako-Saitô-Muñoz-Eaton (WSME) model can be found in earlier works.^43, 44^ The model employs a binary description of residue folded status – *1* for folded and *0* for unfolded – enabling the discretization of protein conformational space into strings of *0*s and *1*s, i.e. every model microstate is effectively represented as a string. Furthermore, specific sequence approximations – single sequence approximation (SSA) wherein only one stretch of folded residues (or *1*s) is allowed, double sequence approximation (DSA) with two stretches of folded residues separated by *0*s (unfolded) are allowed, and DSA allowing for interactions across the folded islands^45^ (DSAw/L) – and a block size of 2 (i.e. two consecutive residues fold-unfold in unison), are considered to reduce the number of microstates for A16 and A749 to 284,198 and 289,122, respectively.^46^ The energy function of the model includes stabilizing van der Waals interactions (with *ξ* as the interaction energy per native contact) from the native structure identified with a 5 Å distance cut-off including nearest heavy atoms (*E_vdW_*), and charge-charge interactions without a distance cut-off employing a Debye-Hückel formalism (*E_elec_*). A temperature-dependent solvation term (Δ*G_solv_*) is also introduced that scales with the number of van der Waals interactions within a microstate and the heat capacity change per native contact (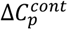). In addition, a large entropic penalty of Δ*S_conf_* is assigned to fix unfolded residues to the folded configuration The thermodynamic parameters that best reproduce the heat capacity curves are: *ξ* of −67.5 ± 2.1 (A749) and −99.8 ± 1.2 J mol^−1^ (A16), Δ*S_conf_* of −14.2 ± 0.79 and −21.5 ± 0.37 J mol^−1^ K^−1^ per residue for all residues other than proline, glycine, and non-helical residues, and 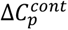 of −0.05 ± 0.02 and −0.63 ± 0.27 J mol^−1^ K^−1^ per native contact. All non-helical and glycine residues are assigned an additional entropic penalty of −6.06 J mol^−1^ K^−1^ per residue, and the entropic cost of fixing proline is set to zero due to its restricted freedom of movement. For A749, the model predicts the first helix to be unfolded or weakly coupled by default, and hence the interactions between H1 and H4 were scaled by a factor of 1.9 to capture experimental trends. To mimic the shape of A749 heat capacity curve in the presence of C18-CoA, the parameters *ξ* and Δ*S_conf_* (of only those residues that bind the ligand) are assigned the values of −67.5 J mol^−1^ and −12.2 J mol^−1^ K^−1^ per residue, respectively. Coupling free energies, free energy profiles and surfaces were generated as described before.^43^

### All-atom molecular dynamics (MD) simulations

Alphafold2 predicted structures of A16 and A749 were placed in a dodecahedral box with a padding distance of 15 Å while maintaining periodic boundary conditions, solvated with TIP3P water molecules (13254 and 12037 water molecules for A16 and A749, respectively), neutralized with ions, and simulated with the AMBER99SB*-ILDN forcefield in GROMACS 2019.6.^47^ The charge-neutralized system was energy minimized using the steepest-descent algorithm before two 500-picosecond relaxation runs - NVT conditions (310 K) with stochastic velocity-rescaling followed by relaxation under NPT conditions (310 K, 1 bar, Parrinello-Rahman barostat). Long-range Coulomb interactions were calculated by employing the particle mesh Ewald (PME) scheme at a grid spacing of 1.2 Å, and a 10 Å cut-off for non-bonded interactions. Equations of motions were solved by employing the leap-frog algorithm with a time-step of 2 femtoseconds. Both the systems were simulated for 5 microseconds and the resulting trajectories analyzed using built-in GROMACS commands. Binding pocket volumes were determined through the standalone software POVME 2.0 by feeding in the trajectory snapshots from MD simulations.^48^ The pocket is constructed by identifying residues within 4 Å of the ligand employing the palmitoyl-CoA bound structure of bovine ACBP (1NVL) as reference.

## Results and Discussion

### Weak unfolding cooperativity in A749

All experiments reported here were carried out in pH 4 and 100 mM ionic strength sodium acetate buffer, conditions at which the unfolding transitions are fully reversible with both proteins exhibiting high solubility. Moreover, little differences in stabilities were observed between pH 4 and pH 7 conditions (Figure S2). Far-ultraviolet circular dichroism (far-UV CD) monitored thermal melting of the two paralogs at 222 nm reveal sigmoidal unfolding profiles with A749 being more stable by 8 K (Figure 1C). At 298 K, this translates to stabilities of 19.8 and 18.1 kJ mol^−1^ for A749 and A16, respectively. Although differences in stabilities between paralogs or orthologs are common observations, the unfolding curve of A749 is significantly broader than that of A16. This difference in cooperativity is quantified by the dimensionless parameter, the Thermodynamic Cooperativity Ratio (TCR), defined as the ratio of the enthalpy of unfolding at the midpoint of A16 over A749 (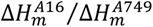 with the Δ*H_m_* values estimated by two-state fits). TCR is a direct measure of sequence- and hence intramolecular interaction-network-driven differences between paralogous proteins, as the overall sequence length and topology are otherwise identical. We obtain a TCR of 1.21 from far-UV CD unfolding curves implying that A16 is 21% more cooperative than A749, despite the lower melting temperature and stability. In addition, the CD signal at 222 nm is significantly weaker (less negative) for A749 compared to A16, suggesting that the native ensemble of A749 is less helical (Figure 1C).

To explore this further, we performed differential scanning calorimetry (DSC) experiments that provide information on the nature of the unfolding transition and probe-independent estimates of cooperativity differences. The resulting absolute heat capacity profile of A749 is strikingly broader than for A16 with a TCR of 1.30 (Figure 2A and Figure S3), and in agreement with estimates from far-UV CD experiments. Furthermore, neither of the profiles can be fit to a two-state model as the folded and unfolded state baselines cross, with the crossing of baselines happening at a lower temperature (320 K) for A749 compared to A16 (335 K), despite the higher *T_m_* of A749 (Figure S3). In other words, neither of the systems can be described by a two-state model and a more complex treatment is required to extract the underlying distribution of states. pfACBPs harbor two conserved tryptophan residues, W60 and W86, with the latter located at the interface of H1 and H4 (Figure 2B). The emission maximum of the tryptophan residues is more red-shifted in A749 (338 nm) compared to A16 (335 nm) even at the lowest temperature (Figure 2C). The thermal unfolding profiles also exhibit larger differences in cooperativity than far-UV CD or DSC, with a TCR of 1.6. In addition, the melting temperatures differ by only 4 K *via* fluorescence (322 K and 326 K for A16 and A749, respectively) while it is 10 K by DSC (322 K and 332 K, respectively). Figure 2D plots the *T_m_* values from different probes highlighting that A16 exhibits a similar melting temperature, while A749 displays a Δ*T_m_* of 6 K with the structural change monitored by tryptophan residues exhibiting the lowest *T_m_*. The inference is that the structural regions around the tryptophan residues melt thermodynamically earlier in A749.

**Figure 2.**
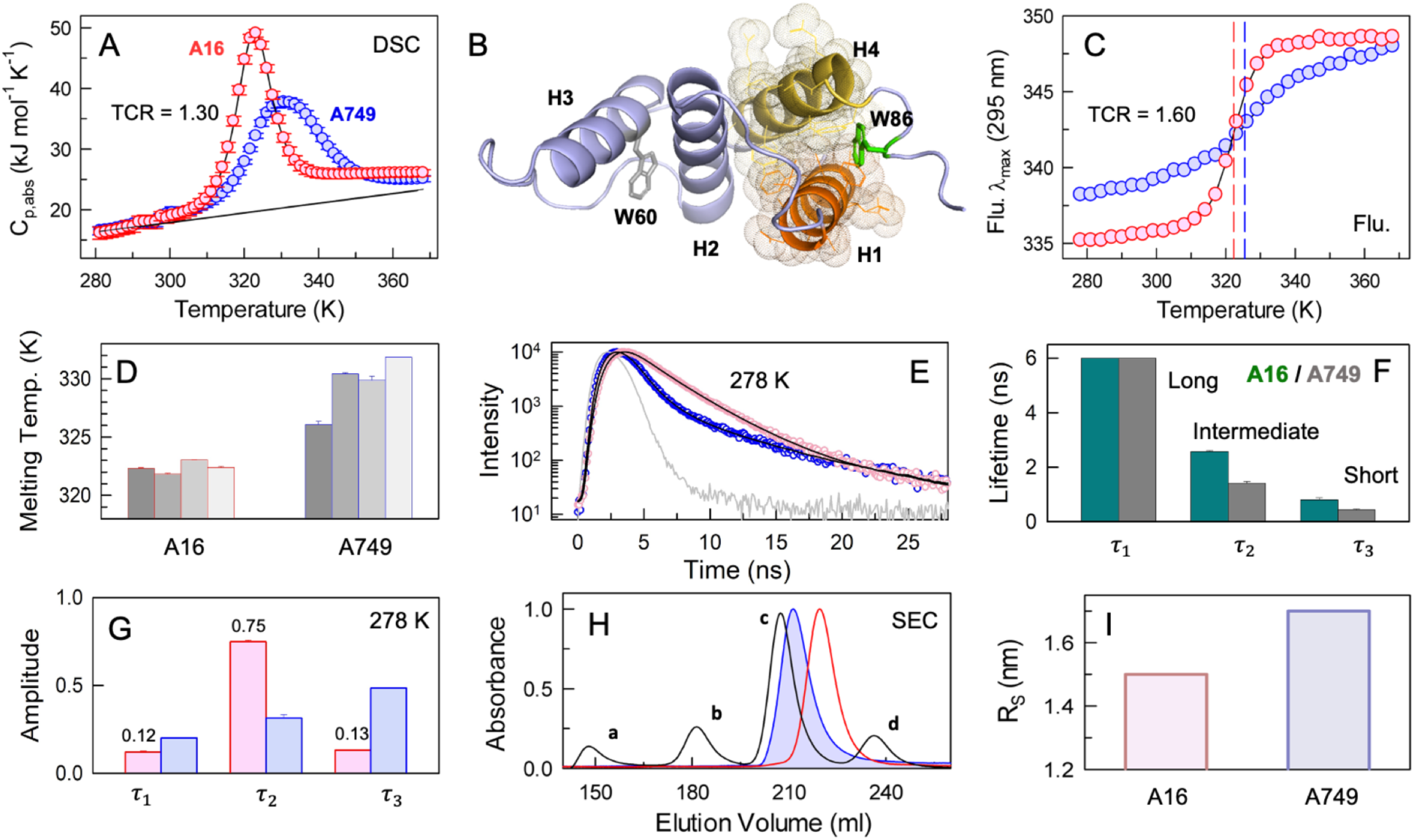
Weak unfolding cooperativity and a broad native ensemble in A749. (A) Absolute heat capacity curves of A16 (red circles) and A749 (blue circles). TCR represents the thermodynamic cooperativity ratio and the black line is the Freire baseline. Note the large difference in cooperativity between the two proteins. (B) Structure of A749 highlighting the relative positioning of helices and the tryptophan residues. The surface representation shows the packing between helices H1 and H4 that is mediated by W86. (C) Fluorescence (Flu.) emission maximum wavelength (*λ_max_*) as a function of temperature on excitation at 295 nm. The vertical dashed lines signal the melting temperatures. (D) Melting temperatures from two-state fits to temperature dependent fluorescence emission maximum, far-UV CD at 222 nm, near-UV CD at 280 nm and DSC (left to right). A749, despite being more stable, displays differences in melting temperatures when monitored by different probes. Error bars are comparable to the line thickness. (E) Fluorescence life-time traces (circles) fit to tri-exponential functions (black) for A16 (magenta) and A749 (blue). Gray trace represents the instrument response function (IRF). (F and G) Extracted life-time components (panel F) and their corresponding amplitudes at 278 K (panel G) following the color code in panel E. Error bars are comparable to the line thickness. (H) Size-exclusion chromatography profiles of A16 (red) and A749 (blue), with the proteins BSA (marked as a), carbonic anhydrase (b), cytochrome C (c) and aprotinin (d) of known Stokes radii as controls to generate a calibration graph. (I) Stokes radius of A16 and A749 extracted from the data shown in panel H.

The contrasting behaviors of tryptophan residues in the paralogs can also be observed from fluorescence life-time (LT) measurements that report on the local electronic environment. Life-time traces at 278 K reveal tri-exponential phases with long, intermediate and short LTs (Figures 2E, 2F) suggestive of folded, partially structured and unfolded-like conformations (monitoring properties of either the backbone or just the indole ring, or a combination of both). Though the LT magnitudes are similar for the paralogs, the amplitudes are distinct and temperature sensitive. Specifically, the longer LT has minimal amplitude for both the paralogs signaling high indole mobility at both 278 K (Figure 2G). However, the intermediate LT has a significantly smaller amplitude for A749 compared to A16 and *vice versa* for the short LT (Figure 2G). We cannot distinguish whether these LTs arise from a single tryptophan or from both, but trends are indicative of larger mobility for the tryptophan residues (or a more fluid structural environment) in A749 compared to A16. Finally, the Stokes radius of A749 is found to be larger than A16 from size-exclusion chromatography (SEC), translating to a 45% higher effective spherical volume for the former at 298 K (Figure 2H, 2I). To summarize, despite displaying a large sequence- and structural similarity to A16, A749 exhibits a more loosely packed core, solvent exposed tryptophan residues with larger mobility, and weaker coupling among its structural elements, manifesting as differences in cooperativity.

### Conformational substates in the native ensemble

We proceeded to acquire ^1^H-^15^N NMR spectra at a range of temperatures to examine conformational differences (if any) at the atomic-level. The spectra recorded between 286-310 K are indicative of folded proteins with well-dispersed resonances (Figure S4). NMR signal assignments for A16 and A749 proteins are currently unavailable. However, we were able to identify the unique peaks observed in the spectral regions characteristic for indole NH tryptophan signals as belonging to the only two tryptophans (W60 and W86) in each of these proteins. We then used site-directed mutagenesis (W60A or W86A) to unambiguously assign those signals to specific tryptophans (Figure S5). A closer look at the tryptophan indole NH region of the NMR spectra reveals that at the lowest temperature of 286 K, both W60 and W86 exhibit a single indole peak as expected of tryptophan residues in a well-folded environment (Figure 3A, 3B). However, on mild temperature increment, W60 starts displaying at least one other minor peak (W60*) that is different from the peak at 286 K. The indole resonances of tryptophan residues in A749 exhibit a similar trend but at a temperature which is 5-6 K higher, corroborating the observations from CD and DSC that A749 is more stable. The major indole peaks (labeled W60 and W86), on the other hand, show an increase in the relative volume as a function of temperature and then decrease sharply, disappearing at ~320 K for A16 and at ~328 K for A749 likely due to exchange broadening (Figure 3C, 3D). The increase in the peak volume is more dramatic for W60 than for W86, and more so in A749 compared to A16. In addition, the relative fraction of the alternate conformational state W60* is larger for A749 and increases with temperature (Figure 3E). Thus, the chemical environment around both W60 and W86 indole side-chains starts changing at significantly earlier temperatures (290-310 K) than the melting temperature, which is at least 10 K higher (red and blue curves in Figure 3C and 3D). In addition, the melting or loss of folded tryptophan resonances (relative to 286 K) occurs thermodynamically earlier for W86 compared to W60.

**Figure 3.**
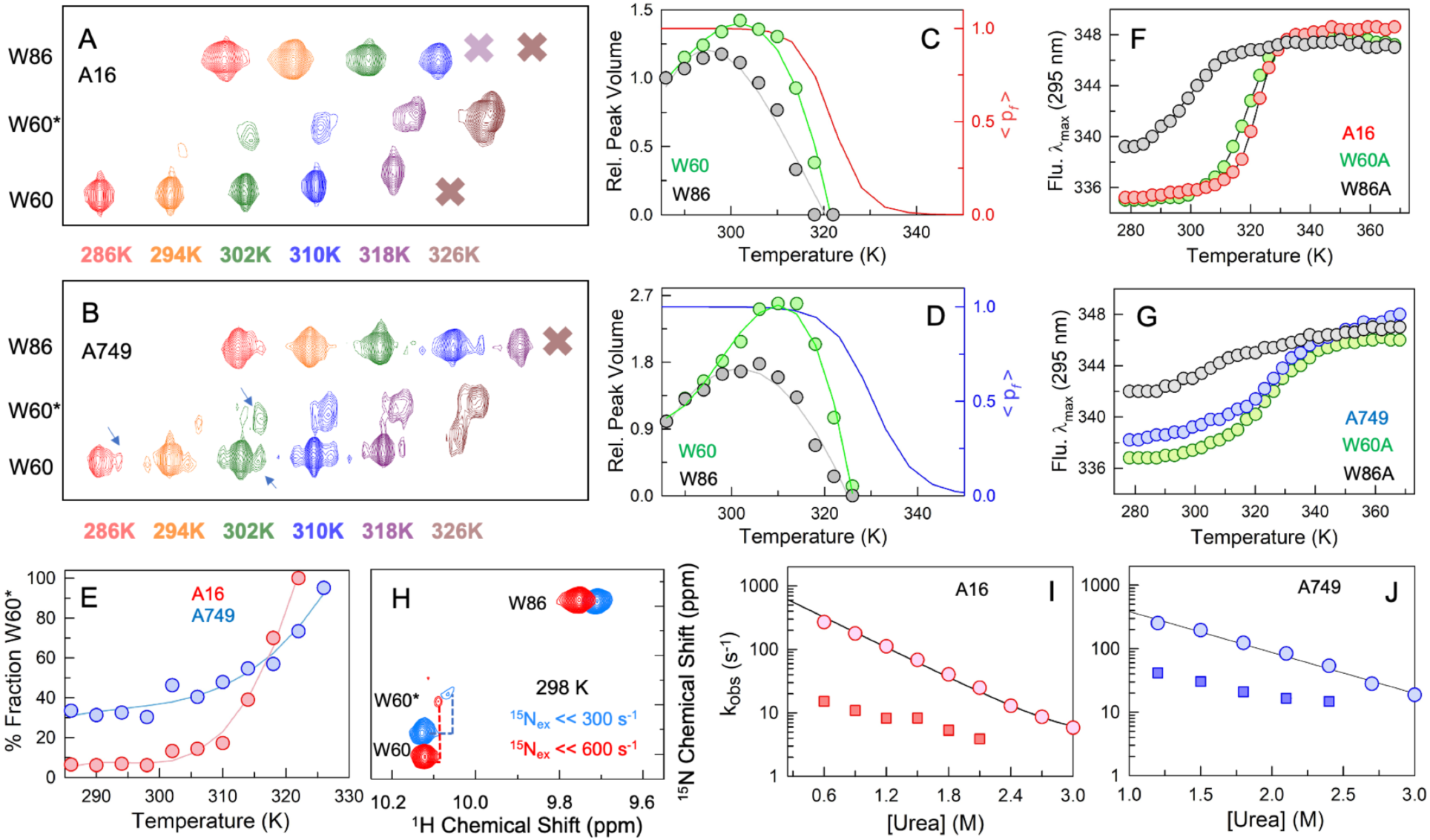
Kinetic complexity and stability determinants. (A and B) A blow-up of the ^1^H-^15^N HMQC spectral region containing W60 and W86 signals, overlaid as a function of temperature for A16 (panel A) and A749 (panel B). The spectra are color-coded by the temperature, and uniformly shifted horizontally for clarity. Note the presence of additional resonances corresponding to W60 (termed W60* and indicated by arrows). (C) Relative peak volumes for W60 and W86 in A16 as a function of temperature employing the peak volume at 286 K as a reference. The red curve signals the folded state probability derived from a two-state fit to the far-UV CD unfolding curve at 222 nm (right axis). (D) Same as panel C, but for A749. (E) The W60* peak volume as a percentage of the total signal for W60 (sum of W60 and W60* peak volumes) is plotted with temperature. (F) Thermal unfolding curves monitored by fluorescence emission maximum for tryptophan mutants of A16. Note that in the W60A mutant, it is the emission from W86 alone that is monitored, and *vice versa*. (G) Same as panel F, but for A749. (H) The spectral region encompassing W60 and W86 signals is shown from ^1^H-^15^N HSQC experiments of A16 (red) and A749 (blue) at 298 K. The highlighted differences (dashed lines) between the ^15^N frequencies of W60 and W60* signals were used to provide an upper limit for the exchange rate. (I and J) Stopped-flow measured relaxation rates constants with circles and squares representing the two relaxation rates observed. The line through circles is shown to guide the eye.

Two observations that stand out from analysis of the NMR spectral are the less stable structural environment around W86 and the presence of multiple resonances for W60 (in both A749 and A16). Since W86 is located at the C-terminal region that is expected to be flexible, it is possible that this residue might not be critical for stability. Structurally, W86 sits right at the interface between helices H1 and H4, appearing to mediate long-range interactions between the two helices. In fact, the A749 W86A mutant is destabilized by 20 K (Δ*T_m_*) compared to the WT protein when monitored by fluorescence, and a similar destabilizing effect is also evident in A16 (gray in Figure 3F, 3G). By contrast, W60A mutations have little relative impact on the overall melting temperature (Δ*T_m_* of 3 K) in both the proteins (green in Figure 3F, 3G). Hence, we conclude that W86 contacts at the H1-H4 interface are critical for the conformational stability of the paralogs. In addition, it is observed that W86 unfolds thermodynamically earlier in WT A749 from fluorescence and NMR probes. Taken together, our experiments establish that the earliest (thermodynamically) structural event in the unfolding of A749 is the undocking of W86 from the H1-H4 interface that is primarily held together by long-range interactions. This event should further destabilize additional regions initiating the unfolding of the protein. We cannot, however, identify which one of the structural elements – H1 or H4 – unfold first, and if A16 follows a similar unfolding behavior (*vide infra*).

The exchange rate (*k_ex_*) between the different resonances of W60 indole NH-group at 298-310 K is estimated from NMR spectra^49^ to be significantly lower than 300 s^−1^ (Figure 3H). These could represent either an exchange between folded and unfolded states of the protein or the dynamic interconversion between various conformational substates within the native ensemble. However, CEST experiments with the saturation time period of 400 ms and ^15^N saturation B_1_ field strengths of 6.5, 12.5 and 25 Hz did not show any discernible dip in intensity for W60/W86 signals when W60* resonance was saturated and *vice versa*. Further, ^15^N transverse relaxation dispersion experiments with CPMG field strengths ranging from 25 to 800 Hz did not yield any measurable exchange contributions to W60/W86 indole relaxation. This implies that the ‘states’ W60 and W60* are in extremely slow exchange (>400 ms or a rate constant of at most 2.5 s^−1^) with each other. While it is established that indole side-chain flips typically take place in the nanosecond timescale, the slow millisecond timescale exchange process in pfACBPs is indicative of a rough native landscape wherein the W60 dynamics is potentially coupled to larger conformational changes involving the protein backbone.

Despite this, stopped-flow kinetic experiments at 285 K reveal bi-exponential relaxation traces at low urea concentrations and single-exponential traces closer to the chemical denaturation midpoint for both the paralogs (Figure 3I, 3J, S6). The faster rate displays a chevron-like behavior (Figure S6) with a folding relaxation rate in water of 1000 s^−1^ at 285 K (Figure 3I, 3J). Moreover, at 298 or 310 K, this rate is expected to increase by a factor of 1.4-1.8 due to a decrease in solvent viscosity relative to 285 K. Therefore, the slower rate that ranges between 4-20 s^−1^ in A16 and 20-50 s^−1^ in A749 represents another kinetics process which is distinct from the W60-W60* exchange.

### Statistical modeling predicts a dynamic native ensemble for A749

In order to quantify folding mechanistic differences and to understand the complex processes observed in equilibrium and kinetics, we resort to a semi-quantitative statistical mechanical treatment of the folding process employing the Wako-Saitô-Muñoz-Eaton (WSME) model (see Methods). A *de novo* prediction of the unfolding thermodynamics (i.e. without calibrating the model parameters against the experimental data) with >280,000 microstates reveals an unfolded H1 for A749 even at the lowest temperatures (Figure S7; note the two-stage unfolding of A749). To reproduce the experimental observations, we iteratively modulate the interaction between H1 and the rest of the structure in A749 alone (by tuning the contact map) and simultaneously attempt to capture the single broad experimental heat capacity profile (see Methods and Figure 4A). The predicted A749 thermogram is still sharper compared to the experimental profile and can be considered to represent a lower limit of the conformational heterogeneity in A749. The sharp heat capacity profile of A16 is however reproduced very well by the model (Figure 4A).

**Figure 4.**
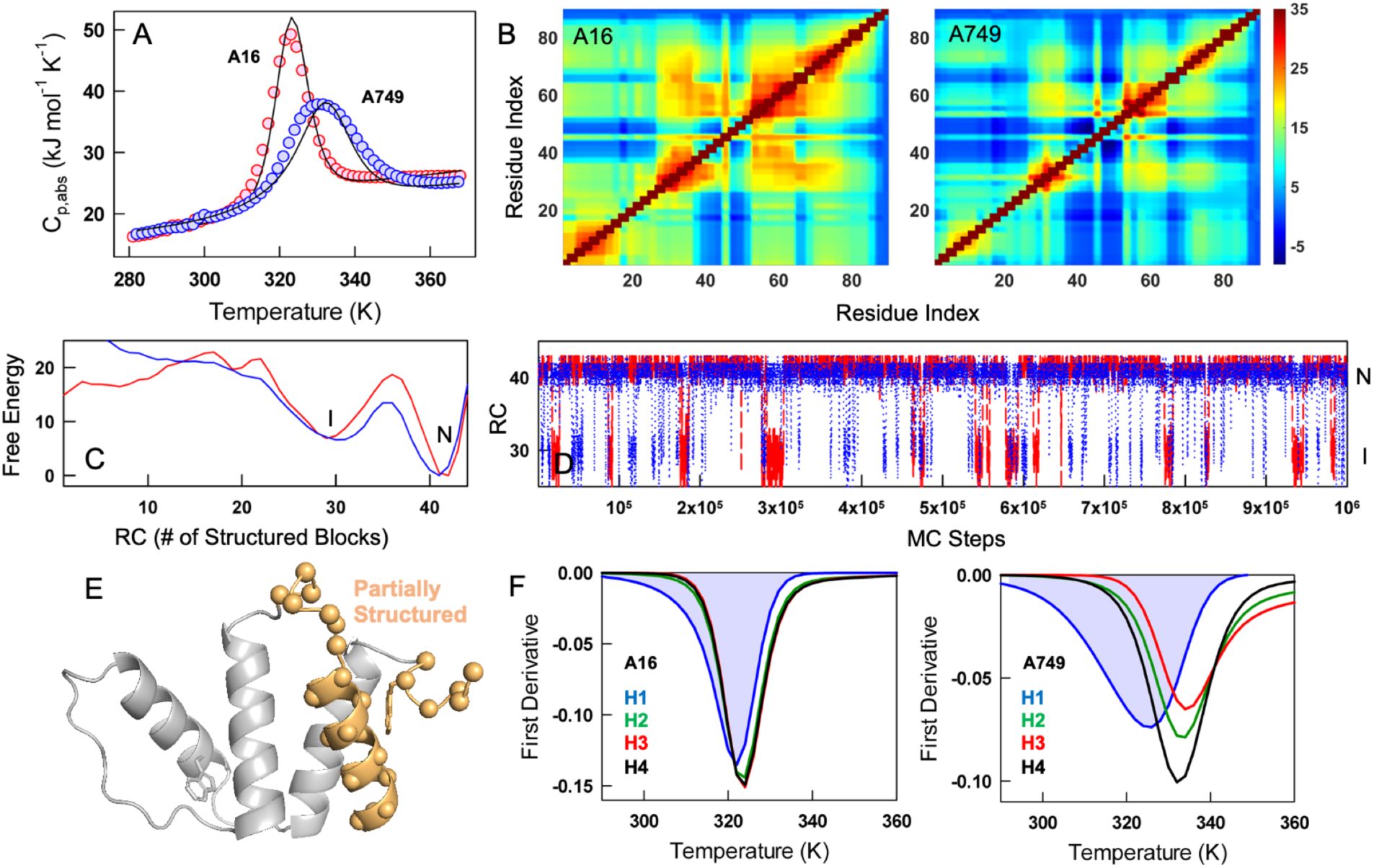
WSME model, coupling patterns and free energy profiles. (A) Fits to the heat capacity profiles from the WSME model (black curves). (B) Effective coupling free-energy matrices at 310 K (in kJ mol^−1^) indicating the weaker coupling in A749 compared to A16. (C) One-dimensional free energy profiles (units of kJ mol^−1^) at 310 K showcasing the presence of an intermediate *I* apart from the native state *N* in both A16 (red) and A749 (blue). (D) Monte-Carlo simulation runs on the profiles shown in panel C highlighting the frequent transitions between *I* and *N* in A749 (blue). (E) Predicted structure of the intermediate whose residues are partially structured or unfolded (orange). (F) Derivatives of predicted unfolding curves from different secondary structure elements for A16 (left panel) and A749 (right panel).

Given the semi-quantitative agreement, we calculate the pair-wise coupling free energies (Δ*G_c_*) between one residue and every other residue that accounts for both the conformational status of a residue in the numerous microstates and the interaction (free-)energies. The coupling map captures the strongly coupled nature of the secondary structure elements and the weak coupling implicit in the long loop connecting H2 and H3 (Figure 4B). A16 displays a relatively stronger coupling tendency among its helices compared to A749, with the differences not localized but spread across the structure. Projecting the conformations onto a reaction coordinate (RC), the number of structured blocks, we find that both the paralogs exhibit a similar feature from the perspective of the one-dimensional free energy profile (FEP) – the presence of an intermediate, *I*, at an RC value of ~30 (Figure 4C). The FEP of A749 displays a smaller barrier between *I* and the native state (*N*) compared to A16. This is more evident from a Monte Carlo simulation on the FEP that highlights the dynamic nature of A749 with the molecule frequently transitioning between *I* and *N* (Figure 4D). A16 is still able to populate the partially structured state *I*, but at a lower frequency and with longer residence times in the *I* state due the larger folding barrier between *I* and *N*.

The structural features of the intermediate are obtained by calculating the mean residue folding probability at the RC value of 30 (Figure S8). The state *I* is predicted to be characterized by partially unfolded H1, loop L1 (connecting H1 and H2), and the C-terminal tail harboring the residue W86 (Figure 4E). Consequently, the unpacking of W86 from the H1-H4 interface could serve as trigger event to decouple H1 from the rest of the structure. This in turn contributes to an earlier (thermodynamically) melting of H1 as observed from first derivatives of helix unfolding curves (i.e. the mean unfolding probability of residues in a given helix; Figure 4F). H1 melts at lower temperatures even in A16 but the stronger thermodynamic coupling ensures that other structural elements follow suit at similar temperatures. On the contrary, the weak thermodynamic coupling in A749 means the protein is able to withstand the melting of H1, and the remaining helices melt at higher temperatures with H2 and H3 unfolding the last. It is pertinent to note that the Δ*T_m_* from the predicted unfolding curves match the experimental range of ~6 K in A749, despite employing DSC curves as the only input. Overall, the WSME model provides an experimentally consistent picture of the folding landscapes of the paralogs and further highlights how small sequence differences can contribute to dramatic differences in the conformational behavior despite a similar unfolding mechanism (*i.e.* through the population of an intermediate).

### Large-scale breathing motions in A749

To further confirm experimental interpretations and WSME model predictions, we performed explicit water all-atom molecular dynamics simulations of the paralogs for five microseconds each. A749 displays higher root mean-squared deviation (RMSD) with respect to the starting structure as a function of simulation time, especially after one microsecond (Figure 5A). The resulting root mean-squared fluctuations (RMSF) are highly contrasting, with the loops of A16 – L1 connecting H1 with H2, and L2 connecting H2 with H3 – exhibiting larger dynamics, while the secondary structure elements are rigid (Figure 5B). On the other hand, the secondary structure elements of A749 are more dynamic while the loops themselves are relatively less flexible, except for the C-terminal tail.

**Figure 5.**
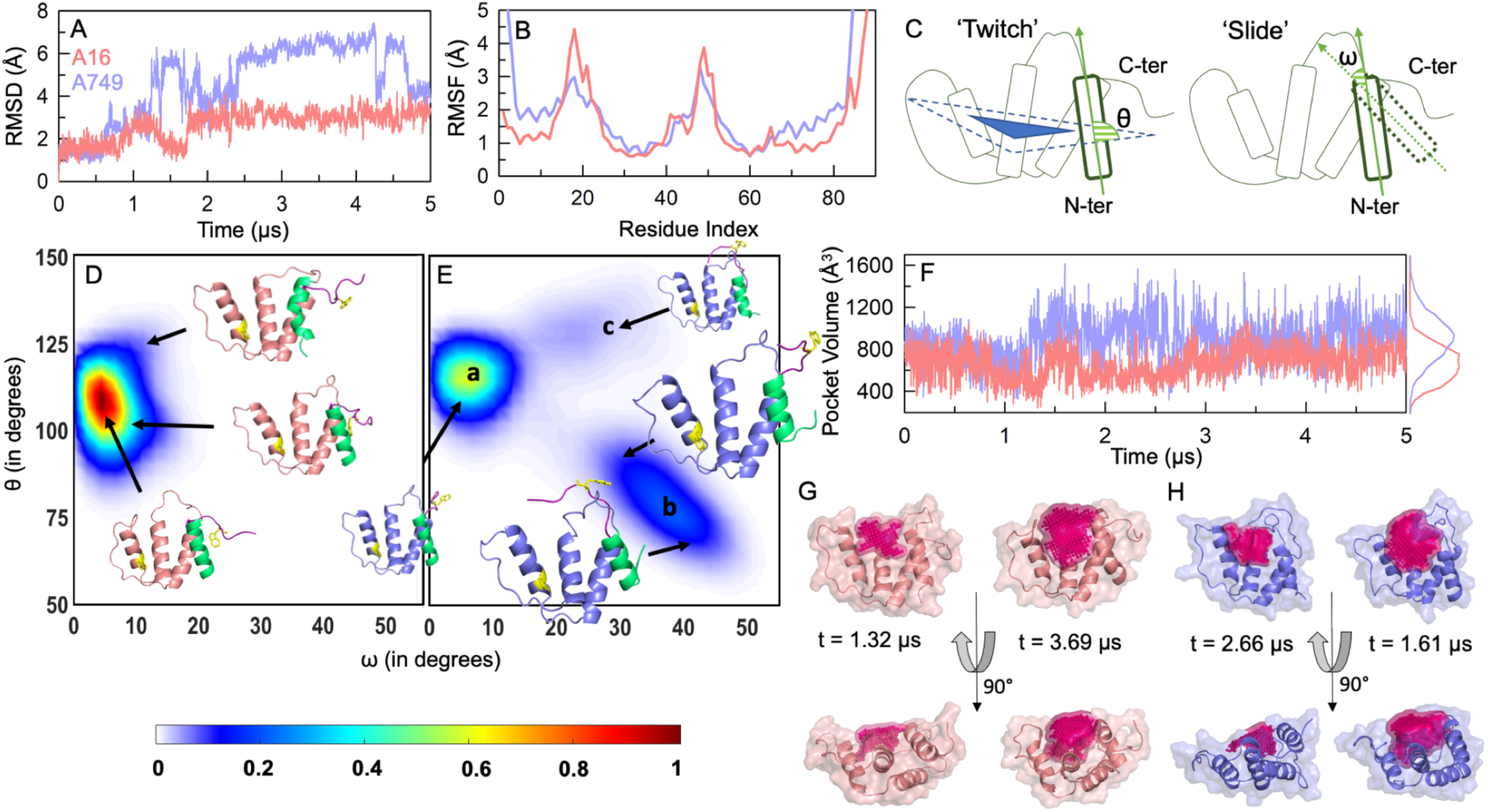
Conformational motions in the native ensemble. (A) Root-mean-square deviation (RMSD) as a function of simulation time. The RMSD starts to change significantly beyond one microsecond simulation time. (B) Root-mean-square fluctuations (RMSF) following the color code in panel A. A749 displays higher RMSF when compared to A16, on average. (C) Cartoon depiction of the structural metrics, twitch and slide, employed to characterize native ensemble dynamics. (D and E) Projection of MD snapshots onto the structural metrics for A16 (panel D) and A749 (panel E). Representative snapshots are highlighted. (F) Binding pocket volume as a function of simulation time following the color code in panel A. The distribution to the right depicts the large binding pocket volume in A749. (G) Specific snapshots that display low (left) and high (right) binding pocket volumes in A16 in two different orientations. (H) Same as panel G, but for A749.

We generate a structural view of the conformational states sampled, by constructing two-dimensional density plots as a function of two variables, *θ* and *ω*, which define the relative orientation of H1 with respect to the plane formed by the other three helices (termed ‘twitch’), and with the starting structure (i.e. at *t* = 0; termed ‘slide’), respectively (Figure 5C). The conformational landscape of A16 is characterized by a relatively rigid H1 orientation, but with minor fluctuations in the C-terminal tail (Figure 5D). A749 accesses a larger conformational space with variations in both the structural metrics (Figure 5E). Specifically, H1 is observed to be quite floppy sampling numerous states, with the majority of the frames exhibiting an unfolded C-terminal tail where W86 is located (labeled ‘*a*’ in Figure 5E). The N-terminal half of H1 is unfolded in some frames and this is observed as shift in both the angles, corresponding to partial melting of structure together with H1 ‘undocking’ from its interactions with H2 and H4 (state ‘*b*’). H1 can also slide along the plane formed by the three helices (H2, H3 and H4) while maintaining its structural integrity, and this can be observed as the minimally populated state ‘*c*’ in Figure 5E. Note that both states *b* and *c* exhibit only ~2.5 turns in the helix H1 (native state is ~3.5 turns).

The consequence of these structural motions is that the dimensions of the acyl-CoA binding pocket (defined by residues spanning H1, L1, H2, and H3) are not rigid, but fluctuate with time (Figure 5F, 5G, 5H). The binding pocket volume of A16 is ~600 Å^3^ on average, but can sample states with a higher pocket volume (~1000 Å^3^) or even a partially closed structure with a pocket volume of just ~400 Å^3^. The binding pocket volume of A749 is consistently higher than that of 16, with a mean value of ~1000 Å^3^, with the protein sampling a spectrum of binding pocket volumes ranging from 400 to 1600 Å^3^, through thermal fluctuations alone. Though the flexibility of L1 and L2 is larger in A16, the arrangement of the side-chains ensures that that the binding pocket remains smaller. This is in contrast to A749 that can dynamically expand or contract the binding pocket by tuning H1 orientation and structure. It thus appears that A749 is more entropically stabilized than A16, a feature that could play an important role in ligand binding.

### Ligand binding induces structural rigidification and conformational restriction

The experiments and simulations discussed above highlight specific differences in the conformational features of the paralogs with the pocket volume being directly linked to the degree of structure in H1 and L1 elements. Does the selection for function, i.e. acyl-CoA binding, contribute to contrasting behaviors? We carried out a series of experiments in the presence of stearoyl-CoA (C18-CoA) to address this question. A DSC scan of A16 at 1:0.2 protein:ligand molar ratio surprisingly reveals only a marginal stabilization of the protein (Figure 6A). A749 is not only more stabilized by the ligand (by 4 K), but a significant increase in cooperativity is observed with a highly asymmetric unfolding transition (Figure 6B). In fact, the sharpness and the maximum C_p_ value of *holo* A749 approaches that of *apo* A16. In other words, the structure of A749 ‘rigidifies’ in the presence of ligand contributing to an increase in cooperativity. While an increase in stability on binding stearoyl CoA is not surprising, the divergent responses of the two proteins are unexpected. Note that proteins are nearly 100% bound by stearoyl-CoA (*K_D_*~100 nM at 298 K for both) under the conditions of experiment (*vide infra*).

**Figure 6.**
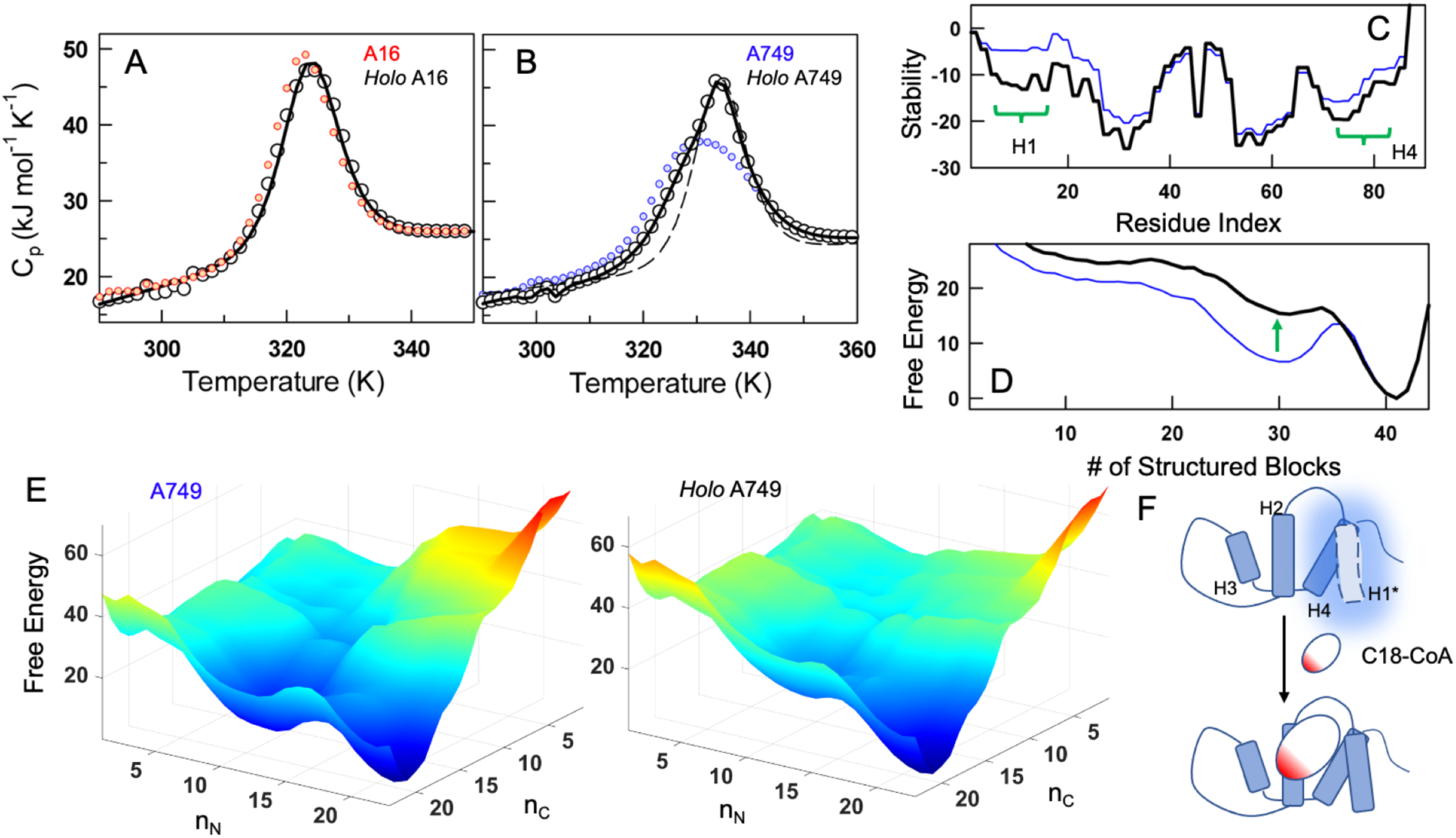
Ligand binding and structural rigidification in A749. (A) Heat capacity profiles of *apo* (red) and *holo* A16 (black circles and black curve) at the protein:ligand molar ratio 1:0.2. (B) Same as panel A, but for A749. The dashed line is the fit from the WSME model wherein ligand-binding residues are rigidified to mimic stabilization conferred by ligand. (C) Residue-level stabilities (in kJ mol^−1^) for A749 at 310 K (more negative signals more stability). An increase in stability is observed for secondary structure elements H1 (by ~10 kJ mol^−1^) and H4 (by ~4 kJ mol^−1^). (D) Free energy profiles of *apo* (blue) and *holo* A749 (black) at 310 K (units in kJ mol^−1^). The population of the intermediate state is nearly fully eliminated on binding the ligand as indicated by the arrow. (E) Free energy surfaces as a function of number of structured blocks in the N- and C-termini (n_N_ and n_C_) display a behavior similar to panel D. Free energy units are in kJ mol^−1^. (F) Cartoons depicting large dynamics of H1 (light blue and dashed boundary) in the *apo* form, which is suppressed in the presence of the ligand.

Since the A749 heat capacity profiles display a dramatic difference between ligand-bound and unbound forms, we quantify the sharpness of the *holo* A749 thermogram employing the WSME model by computationally rigidifying ligand-binding residues (see Methods). At 310 K, the resulting residue level stabilities are enhanced in the *holo* form, and the structural regions that are primarily stabilized involve H1 and L1 that interact with the ligand (Figure 6C). Interestingly, this in turn enhances the stability of H4 (despite this structural element not in direct contact with the ligand), as a consequence of the inter-H1-H4 interactions exhibiting higher stability. From the perspective of the folding free energy profile, the ligand reshapes the profile by reducing the population of the partially structured H1/L1, i.e. the population of the intermediate *I* (Figure 6D).

This feature is also observed in the conformational landscape generated by partitioning the conformational substates into two coordinates involving structured blocks in the N- and C-terminal (Figure 6E). As a consequence, the dynamics of H1 and L1 (which are partially structured in *I*) could be critical for ligand selectivity and affinity. It is, however, important to note that the experimental heat capacity profile of the *holo*-A749 is asymmetric, which is not captured by the WSME model, suggesting that the model predictions are an upper limit to the degree of stabilization of the intermediate state. Effectively, the observations above point to a conformational-selection-like mechanism of ligand binding to A749 (Figure 6F), while there is little evidence for large-scale conformational changes in A16 from DSC.

DSC reports on enthalpic fluctuations with temperature and hence the overall unfolding thermodynamics, which may or may not be sensitive to local and subtle structural changes. It is possible to observe more intricate effects of ligand binding by monitoring local conformational probes. Specifically, fluorescence life-time traces of A16 and A749 in the presence of stearoyl-CoA reveal tri-exponential phases similar to the observations in the absence of the ligand (Figure 7A, 7B). The amplitude of the longer life-time (the 6 nanosecond phase) is enhanced in both paralogs, demonstrating that the torsional rigidity of the tryptophan residues (W60 or W86 or both) increases on binding (Figure 7C, 7D). With increasing temperatures, the amplitude of this phase reduces and matches that of ligand-free proteins at temperatures greater than 295 K. As neither of the tryptophan residues are in direct contact with the ligand, the enhancement of torsional rigidity suggests that the *apo*-forms of the paralogs exhibit larger dynamics in the native ensemble that is dampened on ligand binding, and this holds true for both A16 and A749.

**Figure 7.**
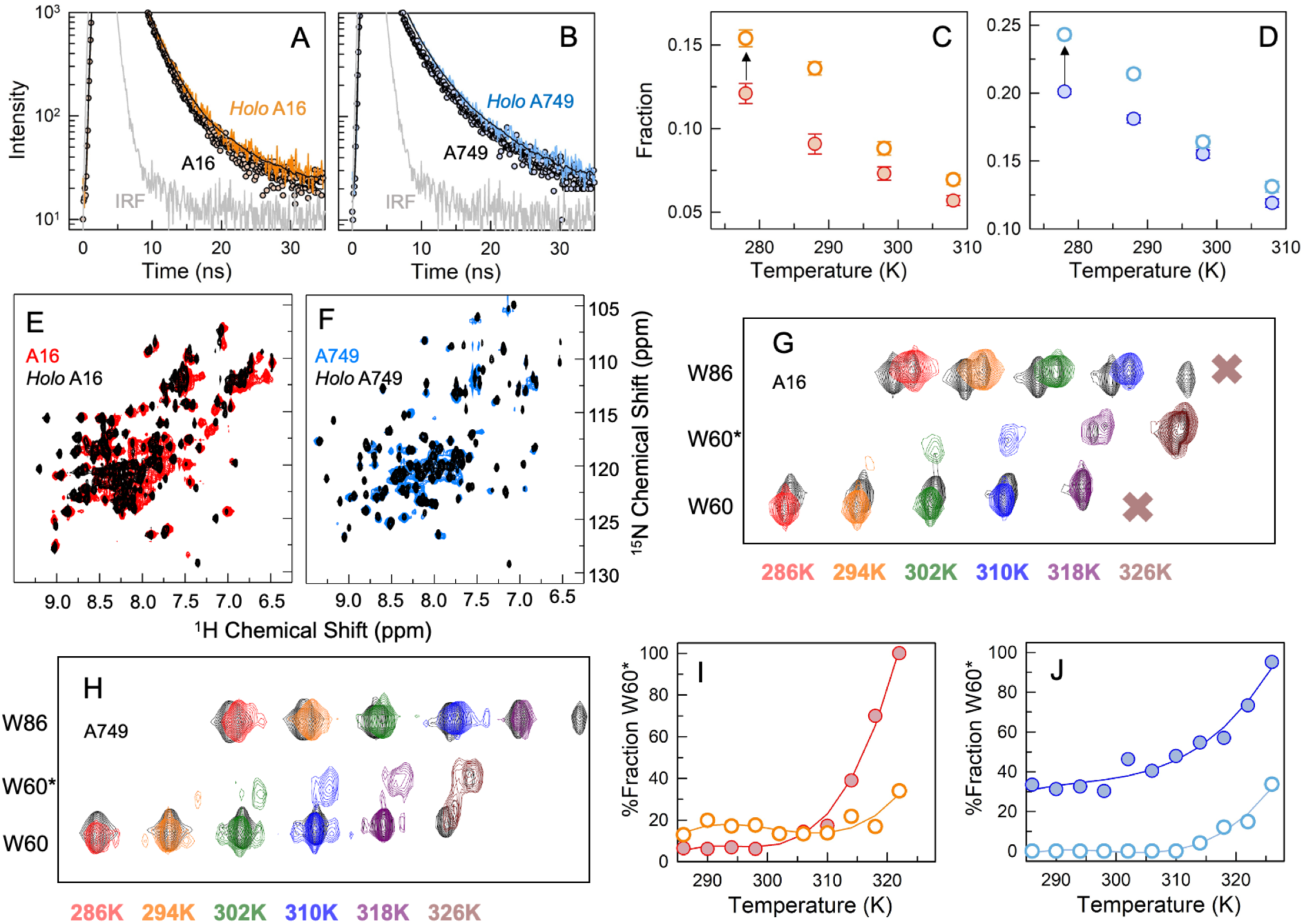
Large-scale structural changes upon binding stearoyl-CoA (C18-CoA). (A and B) Tryptophan fluorescence life-time traces at 278 K for A16 (panel A) and A749 (panel B). In the presence of stearoyl-CoA, the traces (orange and blue) extend to longer times with a higher amplitude. (C and D) The amplitude of the 6 nanosecond component (the longest lifetime; see Fig. 2F) increases in the *holo* form (open circles) relative to the *apo* form (filled circles) for both A16 (panel C) and A749 (panel D). (E and F) ^1^H-^15^N HMQC spectra of A16 (red) and A749 (blue) recorded at 310 K with (black) and without bound ligand. (G and H) Overlaid blow-ups of the spectral region containing W60 and W86 signals recorded at the indicated temperatures for A16 (panel G) and A749 (panel H) with (black) and without bound ligand. The spectra are color-coded by the temperature, and a uniform offset is applied horizontally for clarity. The ligand-bound protein has an increased stability to temperature denaturation as evidenced by the persistent signal for W86 and a decreased population of W60* at high temperatures. A uniform offset is applied horizontally for clarity. (I and J) The percentage fraction of the W60* as a function of temperature for A16 (panel I) and A749 (panel J) in the absence (filled circles) and presence (open circles) of stearoyl-CoA.

The observations above largely hold true in the ^1^H-^15^N NMR spectra comparison of the *apo* and *holo* forms (with stearoyl-CoA) of the paralogs. The peaks are sharper in the *holo* form, but are broadened due to conformational exchange in the *apo* form (Figure 7E, 7F). Signals in the tryptophan indole region of the spectra displays a similar behavior, but with differences. First, the peaks are more stabilized in the *holo* form, as expected (Figure S9). Second, the sharp rise in peak volumes in the temperature range 286 – 310 K is significantly minimized in the ligand-bound A749, while A16 displays only minor differences. This observation is a strong evidence that the large peak volume changes associated with W86 in A749 are related to conformational fluctuations which are suppressed in the ligand-bound state. Third, W86 in *holo* A16 shows two peaks likely due to incomplete saturation under these conditions, while only one is detected at 318 K (Figure 7G, S10); however, W86 in *holo* A749 displays a single peak at all temperatures (Figure 7H, S10). Finally, a more interesting feature is observed for W60, which now exhibits a single resonance corresponding to the major state alone at the lowest temperatures (Figure 7G, 7H, S10). Specifically, the suppression of the minor state (W60*) population is stronger in A749 compared to A16 (Figure 7I, 7J). Taken together, the NMR experiments provide strong evidence to the dampening of large-scale conformational changes in the *holo* (bound) form, in agreement with inferences from scanning calorimetry measurements and fluorescence life-time studies.

### Partial sub-functionalization in A16

It is expected that these differing conformational behaviors would lead to different functional outcomes, i.e. binding to cognate ligands. In fact, it is the functional requirement that determines the extent of conformational excursions, or lack thereof. However, surface plasmon resonance (SPR) based binding affinity measurements reveal little differences between the paralogs when challenged by Lauroyl-CoA (C12-CoA) or Stearoyl-CoA (C18-CoA) at pH 4, and neither of them bind hexanoyl-CoA (C6-CoA) (Figure 8A and Figure S11). Although pH 4 was chosen by us for operational reasons, there is little change in protein stability between pH 4 and 7 (Figure S2). Moreover, the binding surface on the ACBPs is entirely positively charged with the residues K34 and K56 locking the 3’-phosphate of the ribose, apart from K26 and R17 (Figure 8B, 8C). The binding face is also fully conserved with the only difference being R17 in A749 replaced by N17 in A16. Hence the effect of differences in protonation on the binding face would be marginal, if any. The ribose 3’-phosphate in acyl-CoA has a pKa of 6.3,^50, 51^ which would be fully protonated at pH 4, and could therefore contribute to similar binding affinities between the paralogs.

**Figure 8.**
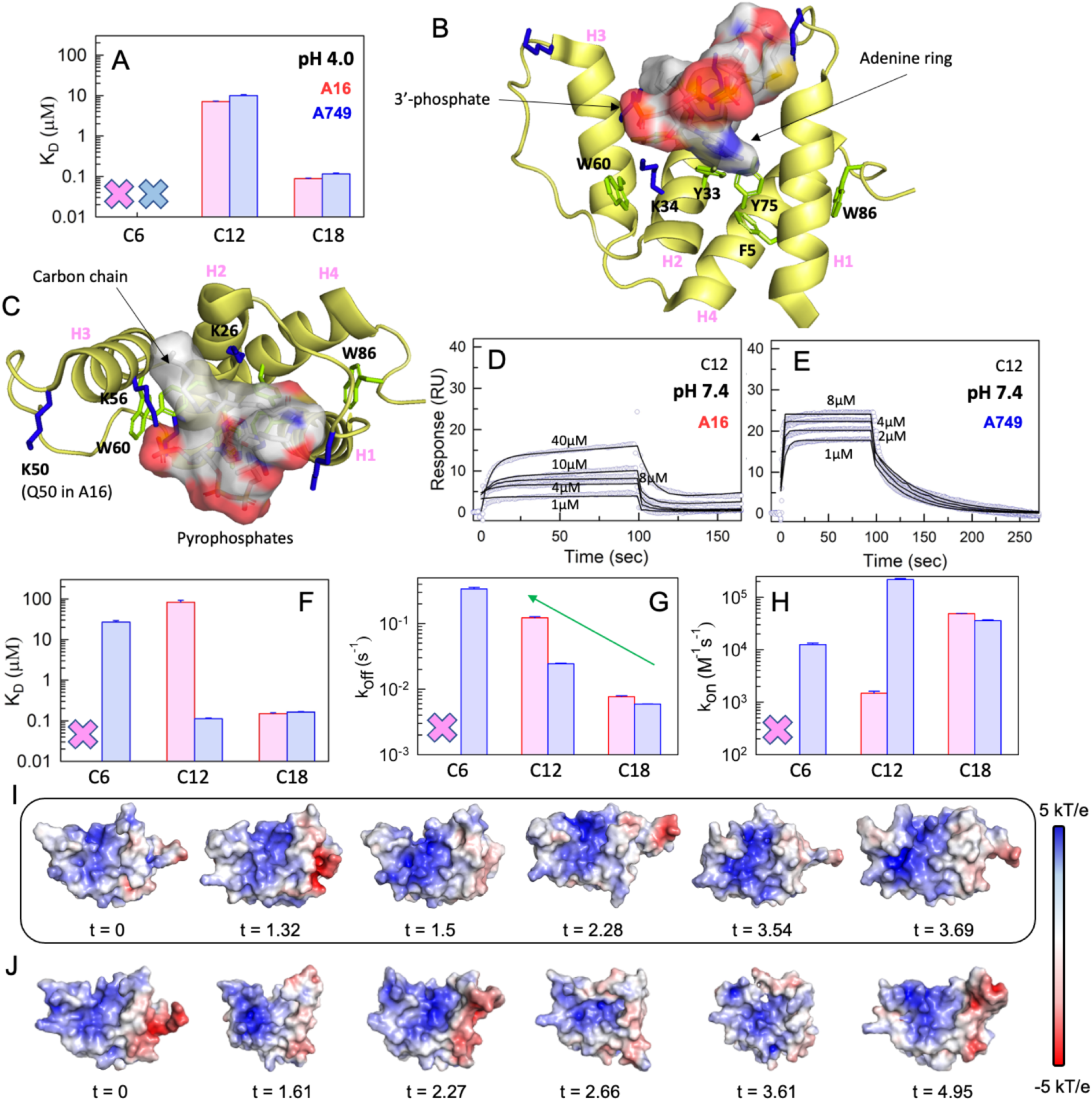
Binding promiscuity and selectivity. (A) Binding affinity of different ligands to A16 (red) or A749 (blue) at pH 4. Crosses indicate very weak or no binding. (B and C) The structure of bACBP bound to palmitoyl-CoA (1NVL) is used to generate a structural model of bound A749, which is shown in two different orientations highlighting the relative positions of charged, tryptophan and tyrosine residues. (D and E) SPR sensograms for A16 and A749 binding C12-CoA at pH 7.4. (F, G and H) Equilibrium dissociation constants (panel F), off-rate (panel G) and on-rate (panel H) for the ligands studied in the current work. Crosses mark weak or no binding or unbinding. (I and J) Electrostatic potential maps at select time points (in microseconds) from MD simulations for A16 (panel I) and A749 (panel J).

We test this expectation by carrying out the SPR experiments in PBS (pH 7.4; Figure 8D, 8E, S12), and find that the dissociation constants of C18-CoA binding to the paralogs are unchanged from pH 4 (Figure 8F). It thus appears that the K34-K56-3’phosphate interaction does not influence the binding equilibrium, and the interactions of the long carbon chain of C18-CoA with the ACBPs likely drive the reaction forward. On the other hand, C12-CoA binds to A749 with a 1000-fold stronger affinity compared to A16. A749 now binds C6-CoA (albeit weakly), while A16 is still unable to bind the shorter carbon chain-length CoA at the higher pH (Figure 8F). Therefore, the differences in protonation of the ribose 3’-phosphate play a critical role in determining the affinity of the short chain fatty-acyl CoAs to ACBPs, similar to earlier observations.^52, 53^ The differences in binding affinities between the paralogs could arise either from the on-rate (*k_on_*; binding-rate) or the off-rate (*k_off_*; dissociation-rate), or from a combination of both. One trend that is obvious is the inverse correlation between the carbon chain-length and the *k_off_*, i.e. shorter chain-lengths exhibit a faster *k_off_* compared to the ligands with longer chain-lengths (Figure 8G). Specifically, the dissociation rate of C12-CoA from A16 is five times faster than from A749, hinting that structural re-arrangements upon binding have significant effects on the off-rate. Being the more rigid of the two, A16 is perhaps unable to re-arrange its interaction network contributing to faster dissociation rates than for A749 which is more accommodative due to its weakly coupled interaction network (Figure 4). In addition, the presence of a more hydrophobic isoleucine at position 52 (that packs against the carbon chain) in A749 compared to alanine at the same position in A16, could contribute to the faster off-rates.

The *k_on_* for A16 exhibits an opposite trend as a function of carbon chain-length, *i.e.* the shorter chain C12-CoA binds nearly 60 times slower than the longer C18-CoA (Figure 8H). For A749, the *k_on_* is maximal for C12-CoA while being 5-20 times slower for C18- and C6-CoAs. The surprising observation here is that the on-rate for C12-CoA binding to A749 is ~200 times faster than that to A16. As the fatty-acyl CoAs are strongly negatively charged due to the presence of 3’-phosphate and pyrophosphates (Figure 8B, 8C), we hypothesize that differences in the positive electrostatic potential on the ligand binding face of the paralogs could fine-tune *k_on_*. This expectation is partially borne out from an electrostatic potential map of ACBPs, with A749 exhibiting a stronger positive potential than A16 in the modeled structure (Figure 8I, 8J at *t* = 0). The differences in electrostatic potential are mediated by two positive charges on the ligand binding face, R17 and K50 in A749 that are replaced with N17 and Q50 in A16. While R17 interacts with the pyrophosphates, K50 does not bind the ligand. However, snapshots from the simulation trajectory reveal that the positive potential on the ligand binding face is quite strong and invariant in A16 (Figure 8I). A749, on the other hand, displays dynamic changes in its shape that in turn alter its surface potential in non-intuitive ways (Figure 8J). It thus appears that a combination of favorable electrostatic potential, conformational selection, and a better structural re-arrangement upon binding could be promoting the relatively rapid binding of C12-CoA to A749.

## Conclusions

The paralogs A16 and A749 as unique systems wherein the functional features – ligand selectivity and affinity - are uniquely tied to the extent of thermodynamic fluctuations in their respective native ensembles. In the conventional ‘Ohno’ model, gene duplication precedes the evolution of new function, with one of the pair now accumulating mutations and thus eventually conferring enhanced fitness.^1^ However, this model is unlikely (as has also been argued by others^10, 54^) as there are combinatorically many more ways to lose function than to gain a precise subset of mutations that confer functionality before being selected out of the population, unless there is some form of dosage selection immediately on gene duplication. In the IAD (innovation, amplification, divergence) model,^54^ innovation (*i.e.* promiscuous activity or binding) occurs first following which gene duplication takes place. One or both the paralogs can now diverge and retain only a subset of the original functional repertoire (exclusive sub-functionalization) or with overlapping functions (partial sub-functionalization). In the current study, A749 exhibits all the hallmarks of an ancestral protein^55–58^ with promiscuous binding, enhanced stability, weak cooperativity, and larger binding pocket volume, and hence ticking off the ‘innovation’ box in the IAD model. It is likely that the A749 gene duplicated, following which mutations accumulated in one or both the genes (divergence). Because long chain fatty acyl-CoAs are critical for membrane biogenesis, the availability of a paralog with exclusive binding to long chain fatty acyl-CoAs (*i.e.* A16) potentially provided a fitness advantage, which was eventually ‘fixed’ in the population along the evolutionary timeline. As with many evolutionary questions, since we are dealing with only the end points of evolutionary trajectories, we can only speculate on the events that could have taken place. Mutational paths from A749 to A16 could be reconstructed (combinations of single- and multiple-point mutations) to better understand the precise molecular mechanism, including further studies on A99, the third paralog.

The partial sub-functionalization in A16, as it is able to interact with only a subset of ligands that A749 binds, arises through a combination of factors, including stronger coupling among its structural elements, reduced dynamics, and time-invariant surface electrostatic potential on the ligand binding face, all of which manifest as sharper unfolding cooperativity and a lower mean binding pocket volume. This effectively means that the native ensemble of A16 is distinct from A749 and is conformationally ‘tuned’ to bind only specific ligands (primarily C18-CoA and very weakly to C12-CoA). Strong binding to C6- or C12-CoA, on the other hand, would require large intrinsic flexibility to wrap the structure around the smaller ligand, which is energetically costly when the helical elements are already strongly coupled (or rigid) in the *apo* form. It is interesting to note that none of the sequence differences between the two paralogs affect the ligand-binding residues, except for asparagine 16 (N16) in the paralog A16. Sequence differences between A16 and A749 are primarily concentrated in H1 and in the C-terminal tail following H4, and hence the long-range interactions between H1 and H4 appear to select, control and gate ligand identity. Remarkably, despite these differences in sequence and thermodynamics, the kinetic complexity is conserved between A16 and A749 over three orders of magnitude – the stopped flow detected relaxation rates of ~1000 s^−1^ (in the absence of urea), the slower rate of 4-50 s^−1^, and the very slow <2.5 s^−1^ from NMR (the W60-W60* interconversion). A similar kinetic complexity is also observed in bovine ACBP, though it has been reported to arise from slow structural acquisition in the unfolded state.^25^

W86 in the C-terminal tail following H4 is unique to the *Plasmodium falciparum* paralogs. Mutations that mimic the undocking of W86, *i.e.* the W86A mutants, dramatically destabilize the folded ensemble, suggestive of W86 acting as a ‘molecular glue’ that stabilizes the H1-H4 interface at lower temperatures. Since the undocking of W86 is the first step in the unfolding process at moderately higher temperatures (>298 K), it is tempting to speculate that this event destabilizes the H1-H4 interface even in the WT proteins, contributing to a highly flexible system with molten-globule-like character that can bind different ligands. W60, the other probe, is not only conserved across *Plasmodium* paralogs, but also across humans and yeast. It is critical for function despite not directly interacting with the ligand. The multiple sub-states monitored by W60 in the *apo* form *via* NMR converge to one in the presence of the ligand, while recovering their multi-resonance character with increasing temperatures in both the proteins. Though we are unable to extricate the precise functional roles of W60 and the conformational origins of multiple resonances, it is clear that the increase in resonance volumes matches the temperatures at which W86 starts undocking from the structure, highlighting long-range effects or cross-talk between the two tryptophan residues.

Though W86 undocks from the native structure thermodynamically earlier in both the proteins (more so for A749), it is in fact the difference in the conformational status of H1 that is quite stark. H1 is shown to undergo large-scale conformational motions including partial unfolding even within the 5 microsecond simulation window in A749. This dynamical character is the likely molecular origin of weak cooperativity in A749 as opposed to A16. In the presence of C18-CoA ligand (which has identical binding affinity to both proteins), A749 alone displays a significantly sharper heat capacity profile which can be construed as rigidification of H1 upon ligand binding. This naturally emerges from WSME model simulations on selectively stabilizing ligand binding residues in A749. It is pertinent to note that this is not a standalone observation in *Plasmodium falciparum* ACBPs. First, Hydrogen-deuterium exchange (HDX) NMR experiments on an equally promiscuous bovine ACBP, a distant evolutionary ortholog, reveal low protection factor for H1 in the *apo* form that switches to higher protection factor in the *holo* form.^21^ Second, WSME model based analysis of bovine ACBP (both *apo* and *holo* forms) predicts a similar flexibility in H1, and a conformational-selection-like ligand-binding mechanism.^26^ Third, electron spin resonance (ESR) spectroscopy of bovine ACBP highlight a distinct increase in H1-H4 interface stability on ligand-binding, apart from the stabilization of binding-site residues in H2 and H3.^59^ Finally, humans harbor a N-terminal truncated isoform of ACBP, hACBP D1, that lacks the entire secondary structure element H1.^60^ This therefore suggests that the structure formed of helices H2-H3-H4 can act as an independent folding unit. Taken together, it is possible that the helix H1 in ACBPs in general acts as a ‘selectivity filter’, exhibiting substantial dynamics in the absence of the ligand while the binding-competent state is arrived at by a conformational selection mechanism. We cannot, of course, rule out an induced-fit mechanism of binding or a combination of both the mechanisms, as we do not have kinetic information on the different steps and the associated fluxes. However, the two ACBP paralogs rigidify globally in the *holo* form, as observed directly in the NMR spectra, lower minor state population and in the higher amplitude of the longer life-time component in time-resolved fluorescence.

In conventional thermodynamic models, a higher stability is expected to arise from a more compact structure with minimal conformational excursions. We see the exact opposite behavior in our work wherein the protein with the higher *T_m_* exhibits larger enthalpic fluctuations with weaker energetic coupling among its structural elements. While stability and *T_m_* values are useful thermodynamic parameters, we emphasize that it is critical to quantify features of the native ensemble, which can be quite counter-intuitive as shown in this work. The differences in the properties of the native ensemble are a consequence of the interaction-network differences and are captured by the WSME model, albeit at a coarse level. Given the large number of orthologs of any given protein family, the model can be employed as a first step to explore the extent to which the ensemble properties are modulated by sequence changes (i.e. the extent of conformational tuning), which could point to critical functional origins. These predictions can be further tested through MD simulations and experiments as we show here, establishing a general framework for understanding folding, dynamics and function.

## Author Information

**Corresponding Authors**

Tel: +91-44-2257 4140

## Acknowledgement

The authors are grateful for the support of the Science and Engineering Research Board (SERB; Department of Science and Technology, India) for the grant CRG/2019/000084 to A. N. N. The authors acknowledge the FIST facility sponsored by the Department of Science and Technology (DST, India) at the Department of Biotechnology, IIT Madras (Chennai, India) for the instrumentation. This work was also supported by the National Institutes of Health grant GM065334 to D.F. and by the Ann G. Wylie Dissertation Fellowship to W.P. NMR experiments were performed on instruments supported in part by the National Science Foundation grant DBI1040158.

## Supporting Information

## Abbreviations

WSME, Wako-Saitô-Muñoz-Eaton; NMR, nuclear magnetic resonance; DSC, differential scanning calorimetry; CD, circular dichroism; HSQC, heteronuclear single-quantum coherence; SOFAST-HMQC, band-selective optimized flip-angle short-transient heteronuclear multiple-quantum coherence; CEST, chemical exchange saturation transfer; CPMG, Carr-Purcell-Meiboom-Gill;

## REFERENCES

(1) Ohno, S. Evolution by Gene Duplication. Springer, New York 1970.

(2) Hughes, A. L. The Evolution of Functionally Novel Proteins after Gene Duplication. Proc. Biol. Sci. 1994, 256 (1346), 119–124. https://doi.org/10.1098/rspb.1994.0058.

(3) Zhang, J. Evolution by Gene Duplication: An Update. Trends Ecol. Evol. 2003, 18, 292–298.

(4) Soria, P. S.; McGary, K. L.; Rokas, A. Functional Divergence for Every Paralog. Mol. Biol. Evol. 2014, 31 (4), 984–992. https://doi.org/10.1093/molbev/msu050.

(5) Kondrashov, F. A.; Rogozin, I. B.; Wolf, Y. I.; Koonin, E. V. Selection in the Evolution of Gene Duplications. Genome Biol. 2002, 3 (2), RESEARCH0008. https://doi.org/10.1186/gb-2002-3-2-research0008.

(6) VanderSluis, B.; Bellay, J.; Musso, G.; Costanzo, M.; Papp, B.; Vizeacoumar, F. J.; Baryshnikova, A.; Andrews, B.; Boone, C.; Myers, C. L. Genetic Interactions Reveal the Evolutionary Trajectories of Duplicate Genes. Mol. Syst. Biol. 2010, 6, 429. https://doi.org/10.1038/msb.2010.82.

(7) Conant, G. C.; Wagner, A. Asymmetric Sequence Divergence of Duplicate Genes. Genome Res. 2003, 13 (9), 2052–2058. https://doi.org/10.1101/gr.1252603.

(8) Force, A.; Lynch, M.; Pickett, F. B.; Amores, A.; Yan, Y. L.; Postlethwait, J. Preservation of Duplicate Genes by Complementary, Degenerative Mutations. Genetics 1999, 151 (4), 1531–1545.

(9) Lynch, M.; Force, A. The Probability of Duplicate Gene Preservation by Subfunctionalization. Genetics 2000, 154 (1), 459–473. https://doi.org/10.1093/genetics/154.1.459.

(10) Copley, S. D. Evolution of New Enzymes by Gene Duplication and Divergence. FEBS J. 2020, 287 (7), 1262–1283. https://doi.org/10.1111/febs.15299.

(11) Yang, G.; Miton, C. M.; Tokuriki, N. A Mechanistic View of Enzyme Evolution. Prot. Sci. 2020, 29 (8), 1724–1747. https://doi.org/10.1002/pro.3901.

(12) Mallik, S.; Tawfik, D. S.; Levy, E. D. How Gene Duplication Diversifies the Landscape of Protein Oligomeric State and Function. Curr. Opin. Genet. Dev. 2022, 76, 101966. https://doi.org/10.1016/j.gde.2022.101966.

(13) Tokuriki, N.; Tawfik, D. S. Protein Dynamism and Evolvability. Science (80-.). 2009, 324 (5924), 203–207. https://doi.org/10.1126/science.1169375.

(14) Hilser, V. J.; Dowdy, D.; Oas, T. G.; Freire, E. The Structural Distribution of Cooperative Interactions in Proteins: Analysis of the Native State Ensemble. Proc. Natl. Acad. Sci. U.S.A. 1998, 95 (17), 9903–9908.

(15) Petrovic, D.; Risso, V. A.; Kamerlin, S. C. L.; Sanchez-Ruiz, J. M. Conformational Dynamics and Enzyme Evolution. J. R. Soc. Interface 2018, 15 (144), 20180330. https://doi.org/10.1098/rsif.2018.0330.

(16) Markin, C. J.; Mokhtari, D. A.; Sunden, F.; Appel, M. J.; Akiva, E.; Longwell, S. A.; Sabatti, C.; Herschlag, D.; Fordyce, P. M. Revealing Enzyme Functional Architecture via High-Throughput Microfluidic Enzyme Kinetics. Science 2021, 373 (6553). https://doi.org/10.1126/science.abf8761.

(17) Yang, G.; Hong, N.; Baier, F.; Jackson, C. J.; Tokuriki, N. Conformational Tinkering Drives Evolution of a Promiscuous Activity through Indirect Mutational Effects. Biochemistry 2016, 55 (32), 4583–4593. https://doi.org/10.1021/acs.biochem.6b00561.

(18) Rajasekaran, N.; Suresh, S.; Gopi, S.; Raman, K.; Naganathan, A. N. A General Mechanism for the Propagation of Mutational Effects in Proteins. Biochemistry 2017, 56, 294–305.

(19) Faergeman, N. J.; Wadum, M.; Feddersen, S.; Burton, M.; Kragelund, B. B.; Knudsen, J. Acyl-CoA Binding Proteins; Structural and Functional Conservation over 2000 MYA. Mol. Cell. Biochem. 2007, 299 (1–2), 55–65. https://doi.org/10.1007/s11010-005-9040-3.

(20) Burton, M.; Rose, T. M.; Faergeman, N. J.; Knudsen, J. Evolution of the Acyl-CoA Binding Protein (ACBP). Biochem. J. 2005, 392, 299–307. https://doi.org/10.1042/bj20050664.

(21) Kragelund, B. B.; Knudsen, J.; Poulsen, F. M. Local Perturbations by Ligand-Binding of Hydrogen-Deuterium Exchange Kinetics in a 4-Helix Bundle Protein, Acyl-Coenzyme-A Binding-Protein (ACBP). J. Mol. Biol. 1995, 250 (5), 695–706. https://doi.org/10.1006/jmbi.1995.0409.

(22) Kragelund, B. B.; Osmark, P.; Neergaard, T. B.; Schiodt, J.; Kristiansen, K.; Knudsen, J.; Poulsen, F. M. The Formation of a Native-like Structure Containing Eight Conserved Hydrophobic Residues Is Rate Limiting in Two-State Protein Folding of ACBP. Nat. Struc. Biol. 1999, 6 (6), 594–601.

(23) Kragelund, B. B.; Poulsen, K.; Andersen, K. V; Baldursson, T.; Kroll, J. B.; Neergard, T. B.; Jepsen, J.; Roepstorff, P.; Kristiansen, K.; Poulsen, F. M.; Knudsen, J. Conserved Residues and Their Role in the Structure, Function, and Stability of Acyl-Coenzyme A Binding Protein. Biochemistry 1999, 38 (8), 2386–2394. https://doi.org/10.1021/bi982427c.

(24) Kragelund, B. B.; Heinemann, B.; Knudsen, J.; Poulsen, F. M. Mapping the Lifetimes of Local Opening Events in a Native State Protein. Prot. Sci. 1998, 7 (11), 2237–2248.

(25) Voelz, V. A.; Jager, M.; Yao, S. H.; Chen, Y. J.; Zhu, L.; Waldauer, S. A.; Bowman, G. R.; Friedrichs, M.; Bakajin, O.; Lapidus, L. J.; Weiss, S.; Pande, V. S. Slow Unfolded-State Structuring in Acyl-CoA Binding Protein Folding Revealed by Simulation and Experiment. J. Am. Chem. Soc. 2012, 134 (30), 12565–12577. https://doi.org/10.1021/ja302528z.

(26) Naganathan, A. N.; Sanchez-Ruiz, J. M.; Munshi, S.; Suresh, S. Are Protein Folding Intermediates the Evolutionary Consequence of Functional Constraints? J. Phys. Chem. B 2015, 119, 1323–1333.

(27) Heidarsson, P. O.; Valpapuram, I.; Camilloni, C.; Imparato, A.; Tiana, G.; Poulsen, F. M.; Kragelund, B. B.; Cecconi, C. A Highly Compliant Protein Native State with a Spontaneous-like Mechanical Unfolding Pathway. J. Am. Chem. Soc. 2012, 134 (41), 17068–17075. https://doi.org/10.1021/ja305862m.

(28) Knudsen, J.; Mandrup, S.; Rasmussen, J. T.; Andreasen, P. H.; Poulsen, F.; Kristiansen, K. The Function of Acyl-CoA-Binding Protein (ACBP)/Diazepam Binding Inhibitor (DBI). Mol. Cell. Biochem. 1993, 123 (1–2), 129–138. https://doi.org/10.1007/BF01076484.

(29) Alquier, T.; Christian-Hinman, C. A.; Alfonso, J.; Færgeman, N. J. From Benzodiazepines to Fatty Acids and beyond: Revisiting the Role of ACBP/DBI. Trends Endocrinol. Metab. 2021, 32 (11), 890–903. https://doi.org/10.1016/j.tem.2021.08.009.

(30) Schroeder, F.; Petrescu, A. D.; Huang, H.; Atshaves, B. P.; McIntosh, A. L.; Martin, G. G.; Hostetler, H. A.; Vespa, A.; Landrock, D.; Landrock, K. K.; Payne, H. R.; Kier, A. B. Role of Fatty Acid Binding Proteins and Long Chain Fatty Acids in Modulating Nuclear Receptors and Gene Transcription. Lipids 2008, 43 (1), 1–17. https://doi.org/10.1007/s11745-007-3111-z.

(31) Hou, N.; Li, S.; Jiang, N.; Piao, X.; Ma, Y.; Liu, S.; Chen, Q. Merozoite Proteins Discovered by QRT-PCR-Based Transcriptome Screening of Plasmodium Falciparum. Front. Cell. Infect. Microbiol. 2021, 11, 777955. https://doi.org/10.3389/fcimb.2021.777955.

(32) Kumar, A.; Ghosh, D. K.; Ali, J.; Ranjan, A. Characterization of Lipid Binding Properties of Plasmodium Falciparum Acyl-Coenzyme A Binding Proteins and Their Competitive Inhibition by Mefloquine. ACS Chem. Biol. 2019, 14 (5), 901–915. https://doi.org/10.1021/acschembio.9b00003.

(33) van Aalten, D. M.; Milne, K. G.; Zou, J. Y.; Kleywegt, G. J.; Bergfors, T.; Ferguson, M. A.; Knudsen, J.; Jones, T. A. Binding Site Differences Revealed by Crystal Structures of Plasmodium Falciparum and Bovine Acyl-CoA Binding Protein. J. Mol. Biol. 2001, 309 (1), 181–192. https://doi.org/10.1006/jmbi.2001.4749.

(34) Jumper, J.; Evans, R.; Pritzel, A.; Green, T.; Figurnov, M.; Ronneberger, O.; Tunyasuvunakool, K.; Bates, R.; Žídek, A.; Potapenko, A.; Bridgland, A.; Meyer, C.; Kohl, S. A. A.; Ballard, A. J.; Cowie, A.; Romera-Paredes, B.; Nikolov, S.; Jain, R.; Adler, J.; Back, T.; Petersen, S.; Reiman, D.; Clancy, E.; Zielinski, M.; Steinegger, M.; Pacholska, M.; Berghammer, T.; Bodenstein, S.; Silver, D.; Vinyals, O.; Senior, A. W.; Kavukcuoglu, K.; Kohli, P.; Hassabis, D. Highly Accurate Protein Structure Prediction with AlphaFold. Nature 2021, 596, 583–589. https://doi.org/10.1038/s41586-021-03819-2.

(35) Mirdita, M.; Schütze, K.; Moriwaki, Y.; Heo, L.; Ovchinnikov, S.; Steinegger, M. ColabFold: Making Protein Folding Accessible to All. Nat. Methods 2022, 19 (6), 679–682. https://doi.org/10.1038/s41592-022-01488-1.

(36) Larkin, M. A.; Blackshields, G.; Brown, N. P.; Chenna, R.; McGettigan, P. A.; McWilliam, H.; Valentin, F.; Wallace, I. M.; Wilm, A.; Lopez, R.; Thompson, J. D.; Gibson, T. J.; Higgins, D. G. Clustal W and Clustal X Version 2.0. Bioinformatics 2007, 23 (21), 2947–2948. https://doi.org/10.1093/bioinformatics/btm404.

(37) Narayan, A.; Gopi, S.; Fushman, D.; Naganathan, A. N. A Binding Cooperativity Switch Driven by Synergistic Structural Swelling of an Osmo-Regulatory Protein Pair. Nat. Commun. 2019, 10 (1), 1995. https://doi.org/10.1038/s41467-019-10002-9.

(38) Guzman-Casado, M.; Parody-Morreale, A.; Robic, S.; Marqusee, S.; Sanchez-Ruiz, J. M. Energetic Evidence for Formation of a PH-Dependent Hydrophobic Cluster in the Denatured State of Thermus Thermophilus Ribonuclease H. J. Mol. Biol. 2003, 329 (4), 731–743.

(39) Schanda, P.; Kupce, E.; Brutscher, B. SOFAST-HMQC Experiments for Recording Two-Dimensional Heteronuclear Correlation Spectra of Proteins within a Few Seconds. J. Biomol. NMR 2005, 33 (4), 199–211. https://doi.org/10.1007/s10858-005-4425-x.

(40) Vallurupalli, P.; Bouvignies, G.; Kay, L. E. Studying “Invisible” Excited Protein States in Slow Exchange with a Major State Conformation. J Am. Chem. Soc. 2012, 134 (19), 8148–8161. https://doi.org/10.1021/ja3001419.

(41) Delaglio, F.; Grzesiek, S.; Vuister, G. W.; Zhu, G.; Pfeifer, J.; Bax, A. NMRPipe: A Multidimensional Spectral Processing System Based on UNIX Pipes. J. Biomol. NMR 1995, 6 (3), 277–293.

(42) Vranken, W. F.; Boucher, W.; Stevens, T. J.; Fogh, R. H.; Pajon, A.; Llinas, M.; Ulrich, E. L.; Markley, J. L.; Ionides, J.; Laue, E. D. The CCPN Data Model for NMR Spectroscopy: Development of a Software Pipeline. Proteins 2005, 59 (4), 687–696. https://doi.org/10.1002/prot.20449.

(43) Naganathan, A. N.; Kannan, A. A Hierarchy of Coupling Free Energies Underlie the Thermodynamic and Functional Architecture of Protein Structures. Curr. Res. Struct. Biol. 2021, 3, 257–267. https://doi.org/10.1016/j.crstbi.2021.09.003.

(44) Anantakrishnan, S.; Naganathan, A. N. Thermodynamic Architecture and Conformational Plasticity of GPCRs. Nat. Commun. 2023, 14 (1), 128. https://doi.org/10.1038/s41467-023-35790-z.

(45) Henry, E. R.; Best, R. B.; Eaton, W. A. Comparing a Simple Theoretical Model for Protein Folding with All-Atom Molecular Dynamics Simulations. Proc. Natl. Acad. Sci. U.S.A. 2013, 110 (44), 17880–17885. https://doi.org/10.1073/pnas.1317105110.

(46) Gopi, S.; Aranganathan, A.; Naganathan, A. N. Thermodynamics and Folding Landscapes of Large Proteins from a Statistical Mechanical Model. Curr. Res. Struct. Biol. 2019, 1, 6– 12.

(47) Pronk, S.; Pall, S.; Schulz, R.; Larsson, P.; Bjelkmar, P.; Apostolov, R.; Shirts, M. R.; Smith, J. C.; Kasson, P. M.; van der Spoel, D.; Hess, B.; Lindahl, E. GROMACS 4.5: A High-Throughput and Highly Parallel Open Source Molecular Simulation Toolkit. Bioinformatics 2013, 29, 845–854.

(48) Durrant, J. D.; Votapka, L.; Sørensen, J.; Amaro, R. E. POVME 2.0: An Enhanced Tool for Determining Pocket Shape and Volume Characteristics. J. Chem. Inf. Model. 2014, 10 (11), 5047–5056. https://doi.org/10.1021/ct500381c.

(49) Narayan, A.; Campos, L. A.; Bhatia, S.; Fushman, D.; Naganathan, A. N. Graded Structural Polymorphism in a Bacterial Thermosensor Protein. J. Am. Chem. Soc. 2017, 139, 792–802.

(50) Alberty, R. A.; Smith, R. M.; Bock, R. M. The Apparent Ionization Constants of the Adenosinephosphates and Related Compounds. J. Biol. Chem. 1951, 193 (1), 425–434.

(51) Keire, D. A.; Robert, J. M.; Rabenstein, D. L. Microscopic Protonation Equilibria and Solution Conformations of Coenzyme A and Coenzyme A Disulfides. J. Org. Chem. 1992, 57, 4427–4431.

(52) Kragelund, B. B.; Andersen, K. V; Madsen, J. C.; Knudsen, J.; Poulsen, F. M. 3-DIMENSIONAL STRUCTURE OF THE COMPLEX BETWEEN ACYL-COENZYME-A BINDING-PROTEIN AND PALMITOYL-COENZYME-A. J. Mol. Biol. 1993, 230 (4), 1260–1277. https://doi.org/10.1006/jmbi.1993.1240.

(53) Faergeman, N. J.; Sigurskjold, B. W.; Kragelund, B. B.; Andersen, K. V; Knudsen, J. Thermodynamics of Ligand Binding to Acyl-Coenzyme A Binding Protein Studied by Titration Calorimetry. Biochemistry 1996, 35, 14118–14126.

(54) Bergthorsson, U.; Andersson, D. I.; Roth, J. R. Ohno’s Dilemma: Evolution of New Genes under Continuous Selection. Proc. Natl. Acad. Sci. U. S. A. 2007, 104 (43), 17004–17009. https://doi.org/10.1073/pnas.0707158104.

(55) Ingles-Prieto, A.; Ibarra-Molero, B.; Delgado-Delgado, A.; Perez-Jimenez, R.; Fernandez, J. M.; Gaucher, E. A.; Sanchez-Ruiz, J. M.; Gavira, J. A. Conservation of Protein Structure over Four Billion Years. Structure 2013, 21 (9), 1690–1697. https://doi.org/10.1016/j.str.2013.06.020.

(56) Zou, T.; Risso, V. A.; Gavira, J. A.; Sanchez-Ruiz, J. M.; Ozkan, S. B. Evolution of Conformational Dynamics Determines the Conversion of a Promiscuous Generalist into a Specialist Enzyme. Mol. Biol. Evol. 2015, 32 (1), 132–143. https://doi.org/10.1093/molbev/msu281.

(57) Risso, V. A.; Sanchez-Ruiz, J. M.; Ozkan, S. B. Biotechnological and Protein-Engineering Implications of Ancestral Protein Resurrection. Curr. Opin. Struct. Biol. 2018, 51, 106–115. https://doi.org/10.1016/j.sbi.2018.02.007.

(58) Modi, T.; Huihui, J.; Ghosh, K.; Ozkan, S. B. Ancient Thioredoxins Evolved to Modern-Day Stability-Function Requirement by Altering Native State Ensemble. Philos. Trans. R. Soc. Lond. B Biol. Sci. 2018, 373 (1749). https://doi.org/10.1098/rstb.2017.0184.

(59) Hung, C.; Lee, S. W.; Chiang, Y. Local Structural Stability of the Acyl-Coenzyme A Binding Protein by ESR Spectroscopy. Appl. Mag. Res. 2023, 54, 107–118.

(60) Nitz, I.; Kruse, M.-L.; Klapper, M.; Döring, F. Specific Regulation of Low-Abundance Transcript Variants Encoding Human Acyl-CoA Binding Protein (ACBP) Isoforms. J. Cell. Mol. Med. 2011, 15 (4), 909–927. https://doi.org/10.1111/j.1582-4934.2010.01055.x.

