## Supporting Information for "Conformational Tuning Shapes the Balance between Functional Promiscuity and Specialization in Paralogous *Plasmodium* Acyl-CoA Binding Proteins"

AUTHOR INFORMATION

**Corresponding Author**

[‡]Equal contribution

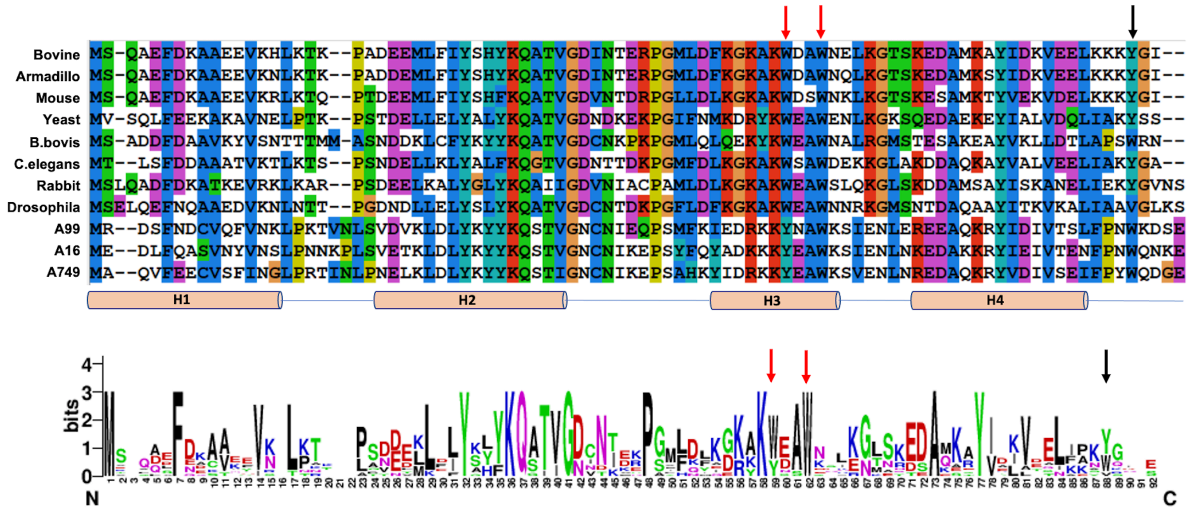


**Figure S1** Sequence alignment (top panel) and WebLogo[1] (bottom panel) of select ACBP paralogs and orthologs. The last three sequences in the top panel represent the *Plasmodium falciparum* paralogs. The arrows indicate the location of conserved aromatic residues. Note that position 86 is predominantly occupied by aromatic residues with tryptophan being conserved across the Plasmodium paralogs. *Babesis bovis* also harbors a tryptophan at position 86 while other eukaryotes do not.

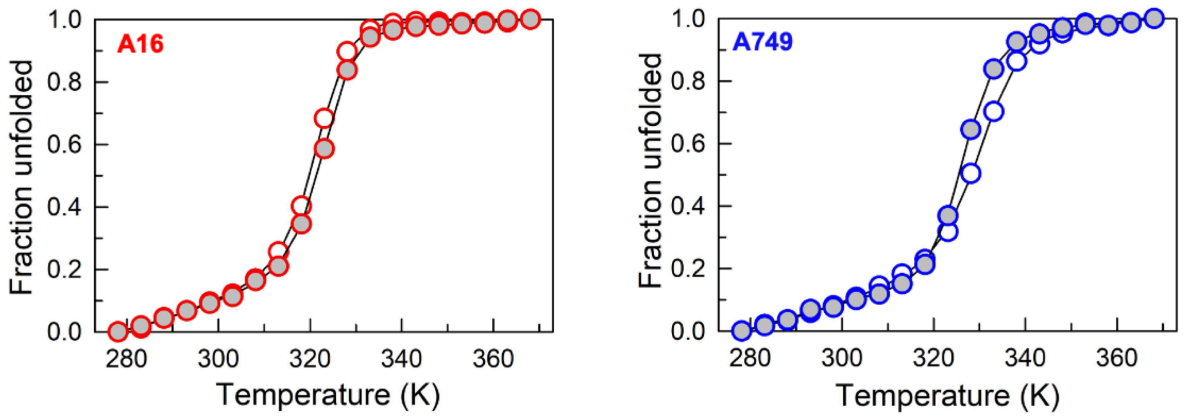


**Figure S2** Comparison of normalized thermal unfolding curves monitored via far-UV CD at 222 nm at pH 7 (filled circles) and pH 4 (open circles).

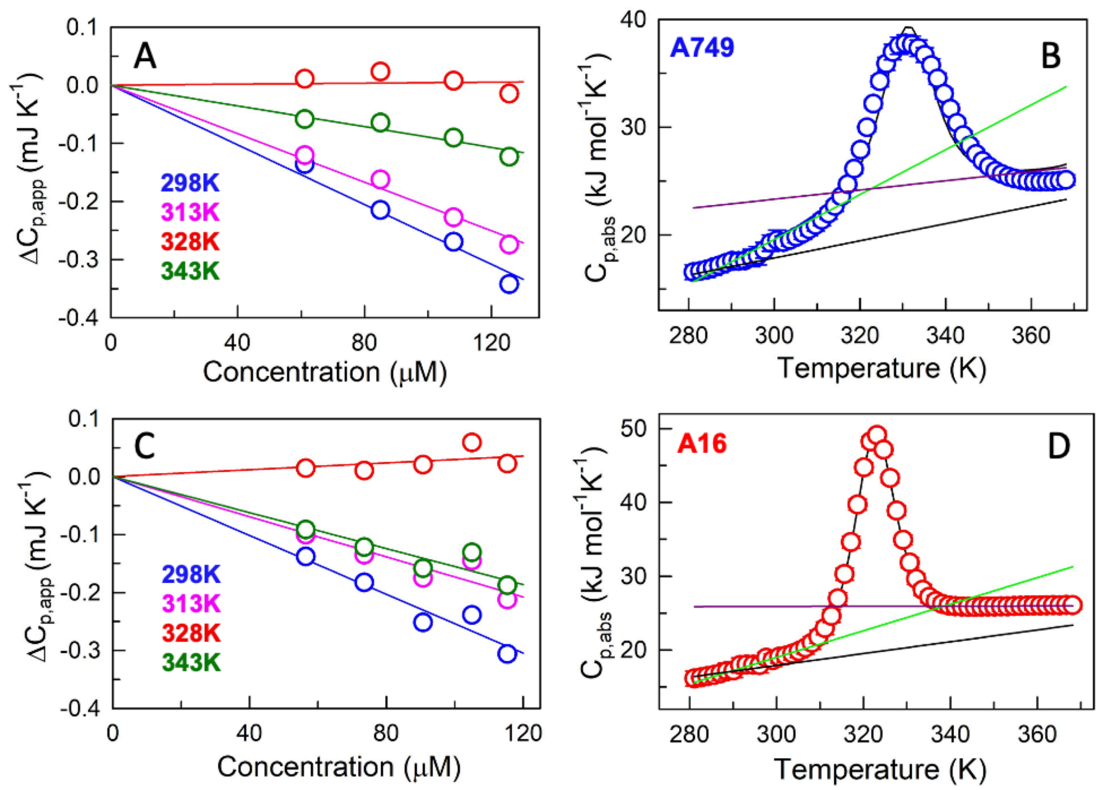


**Figure S3** (A and C) Apparent heat capacities at select temperatures and the range of concentrations studied in this work for A749 (panel A) and A16 (panel C). The slope of the best fit line carries information on the absolute heat capacity.[2] (B and D) Absolute heat capacity profiles of the paralogs fit to a two-state model (curves through the points). Green and purple represent the folded and unfolded state baselines, respectively, that cross within the experimental temperature range. Black line represents the Freire baseline.

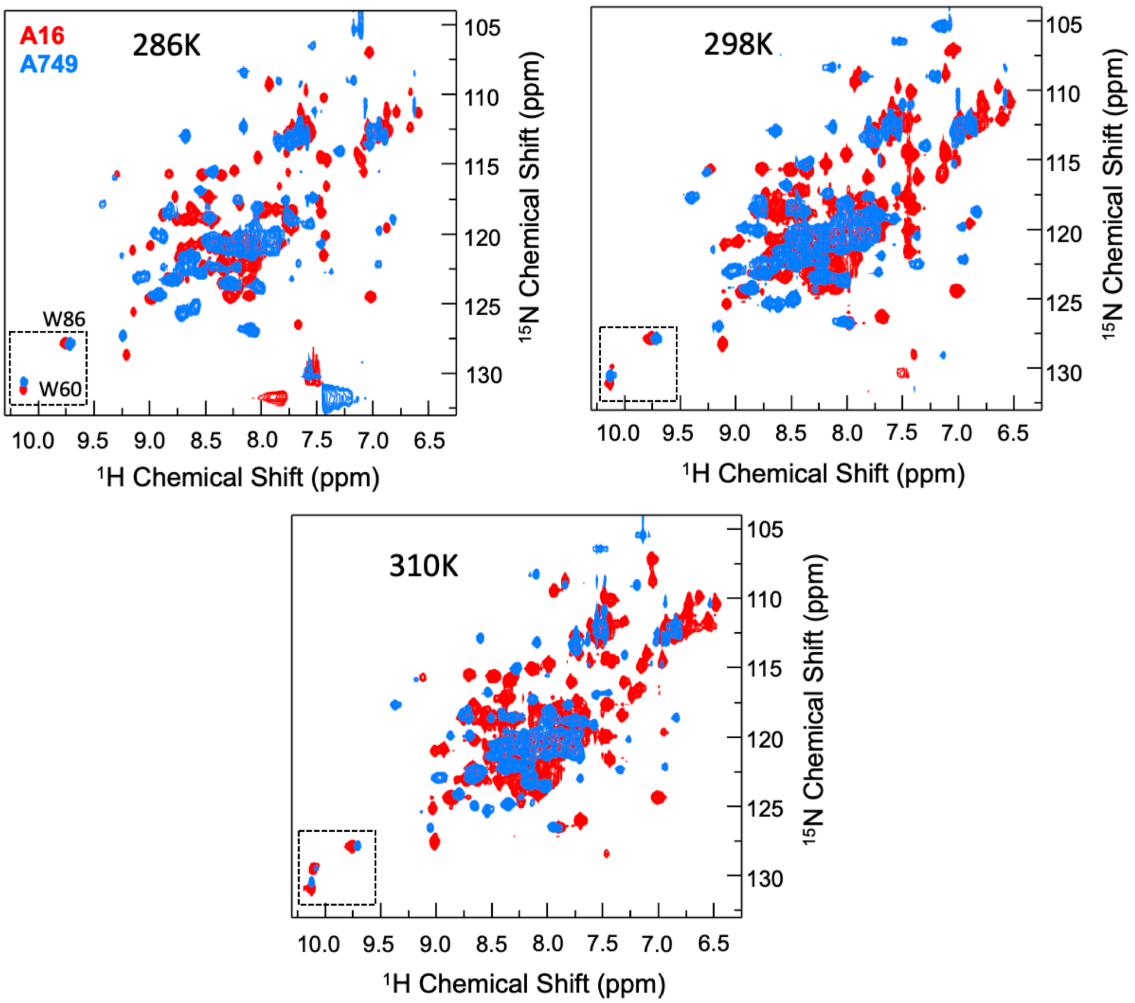


**Figure S4** Overlay of $^{1}$H-$^{15}$N SOFAST-HMQC spectra of A749 (blue) onto A16 (red) recorded at three temperatures. The spectral region containing indole signals of W60 and W86 is indicated by the dashed box. A blow-up of this region is shown in Figure 3A, 3B (main text).

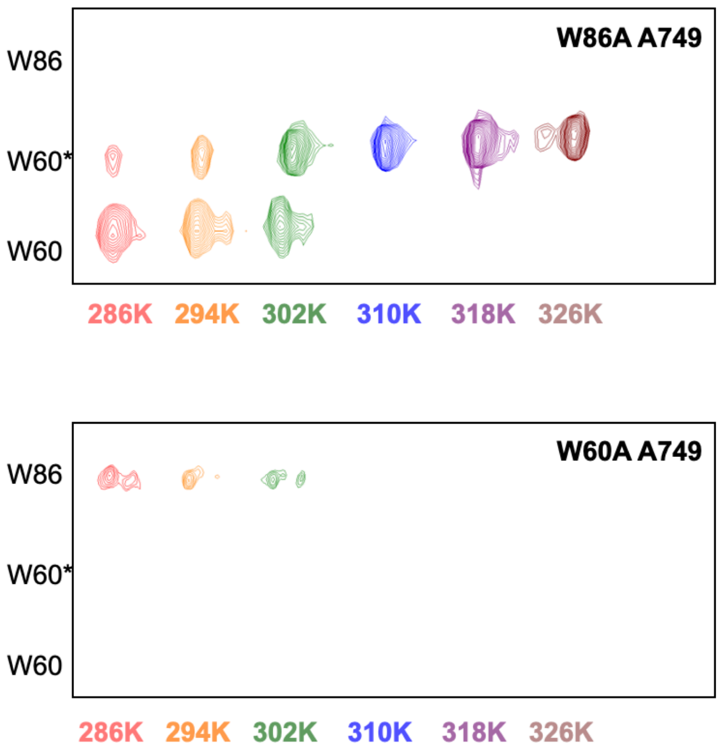


**Figure S5** A blow-up of the tryptophan indole NH region of the NMR spectra for the A749 mutants W86A (top) and W60A (bottom). It can be seen that both W60 and W60* peaks are observable for W86A mutant, but absent in the W60A mutant.

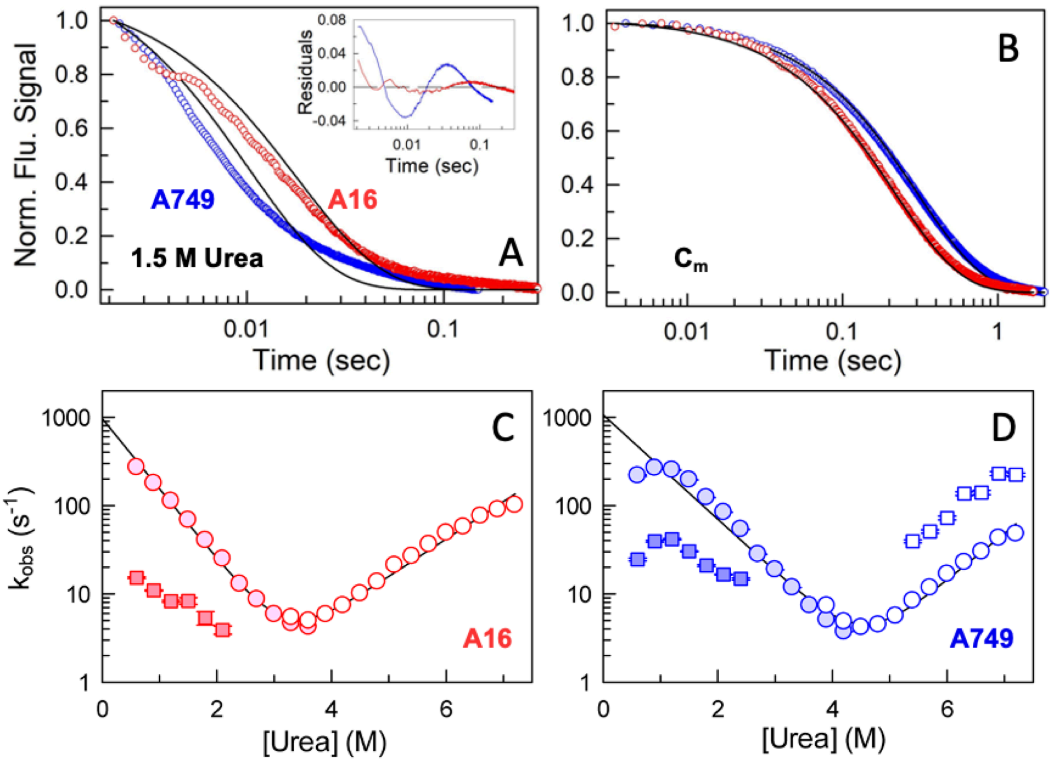


**Figure S6** (A) Stopped-flow relaxation traces at 1.5 M urea (circles) fit to a single-exponential function (black). The resulting residuals are shown in the inset. (B) Relaxation traces at the chemical denaturation midpoint ($C_m$; circles) fit to a single-exponential function (black). (C and D) Folding relaxation rates (filled circles) and unfolding relaxation rates (open circles) that exhibit a Chevron-like behavior. The black curve is a two-state model fit. Both paralogs also exhibit a slower relaxation rate (filled squares) under refolding conditions, while A749 additionally exhibits a faster phase under unfolding conditions (open squares).

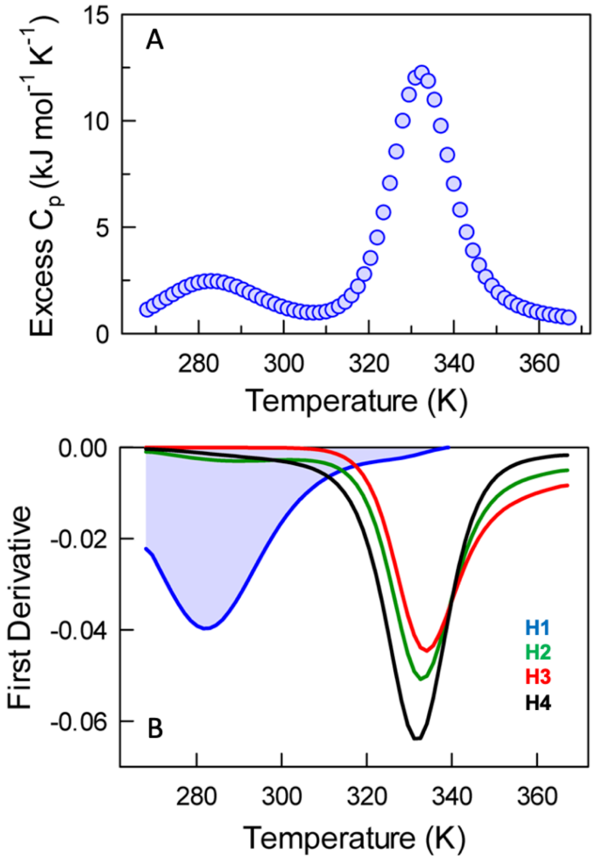


**Figure S7** (A) Predicted excess heat capacity curve of A749 from the bWSME model[3,4] without modulation of contact map and employing identical parameters as noted in the Methods section. (B) First derivatives of the mean residue folding probabilities of various secondary structure elements. Note that helix H1 melts earlier, and this shows up as a bump at ~280 K in the heat capacity profile in panel A.

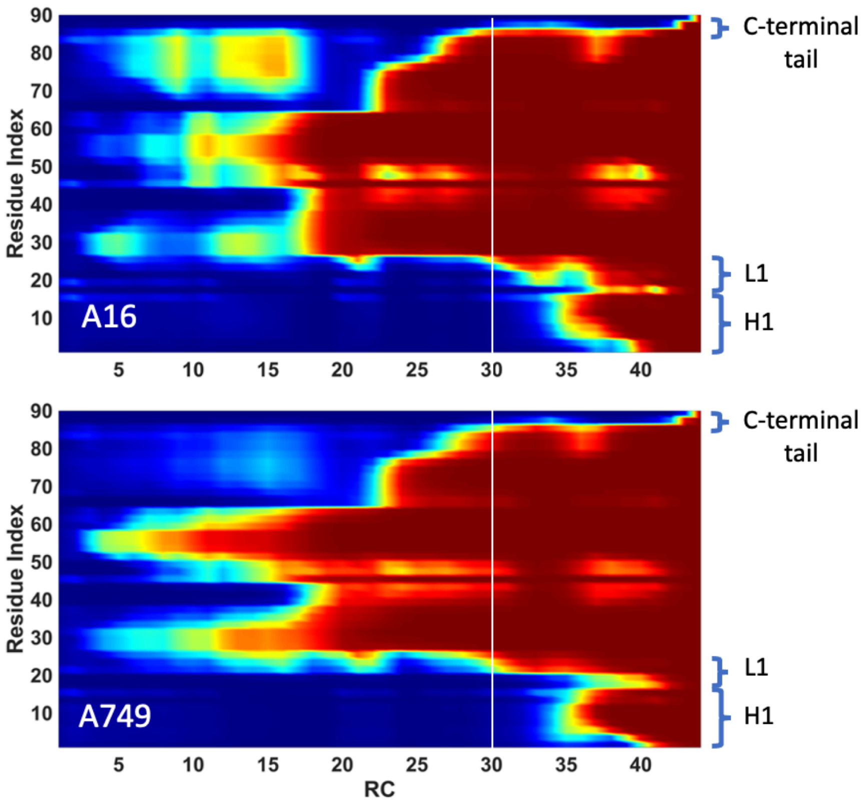


**Figure S8** Plots of probabilities color-coded in the spectral scale from fully unfolded (blue and zero probability) to fully folded (red and probability of 1), as a function of the reaction coordinate (RC; number of structured blocks) for A16 and A749 predicted by the bWSME model. Note that at the RC value of 30 (corresponding to the intermediate in free-energy profiles shown in Figure 4C of the main text), the first 20-25 residues (corresponding to H1 and L1) and the last few residues (C-terminal tail) are unfolded (blue) in both the proteins.

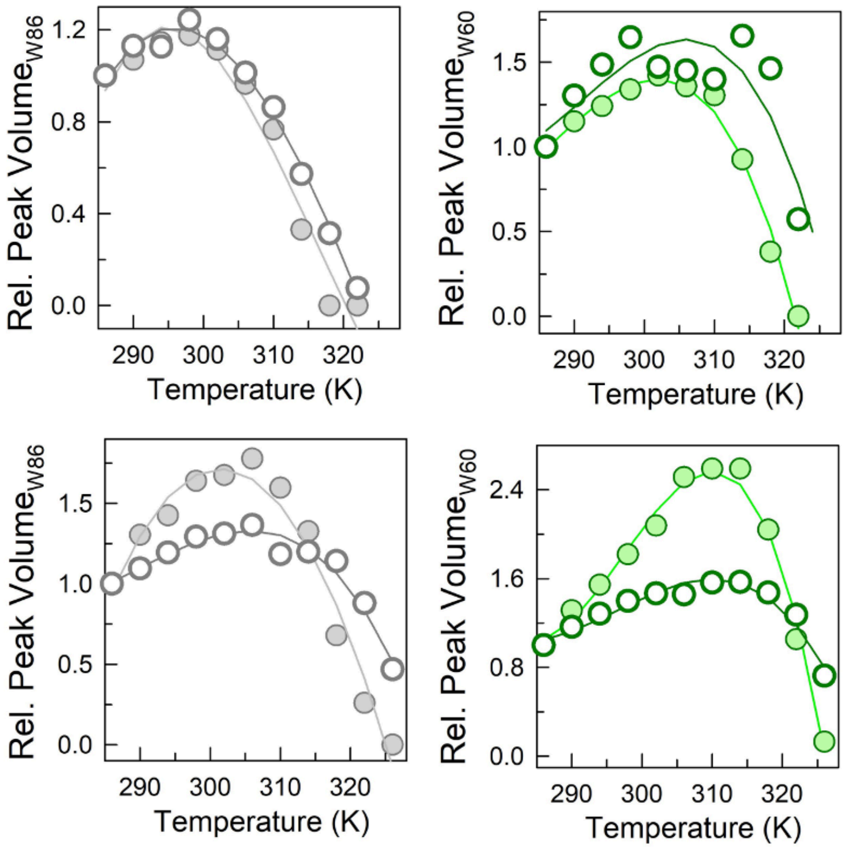


**Figure S9** Relative NMR peak volumes of W86 (left column) and W60 (right column) for A16 (top row) and A749 (bottom row), respectively. Open and filled circles represent bound (*holo*) and ligand-free (*apo*) forms, respectively. Experiments were carried out in the presence of stearoyl-CoA (C18-CoA).

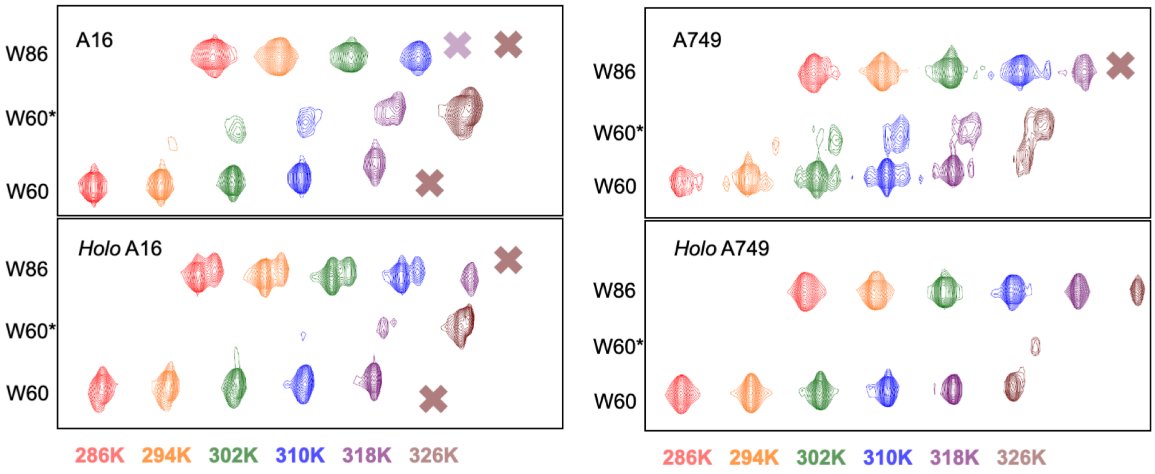


**Figure S10** Effect of ligand binding on the tryptophan indole NH region of the NMR spectra. In Figure 7G, 7H of the main text, the same plots are displayed but with the *holo* form shown in black and superimposed onto the *apo* form (top row).

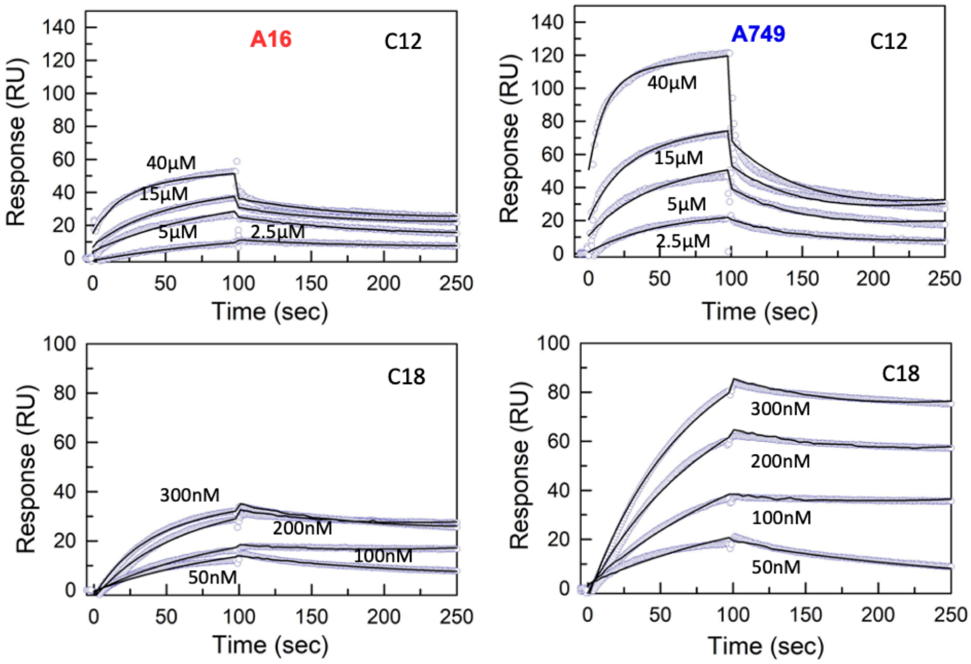


**Figure S11** Surface plasmon resonance (SPR) sensograms of different ligands binding to and unbinding from A16 (left column) and A749 (right column) at pH 4. Note the extremely slow unbinding of C18 from both the paralogs. Black lines passing through the data represent global fit from a 1:1 binding model.

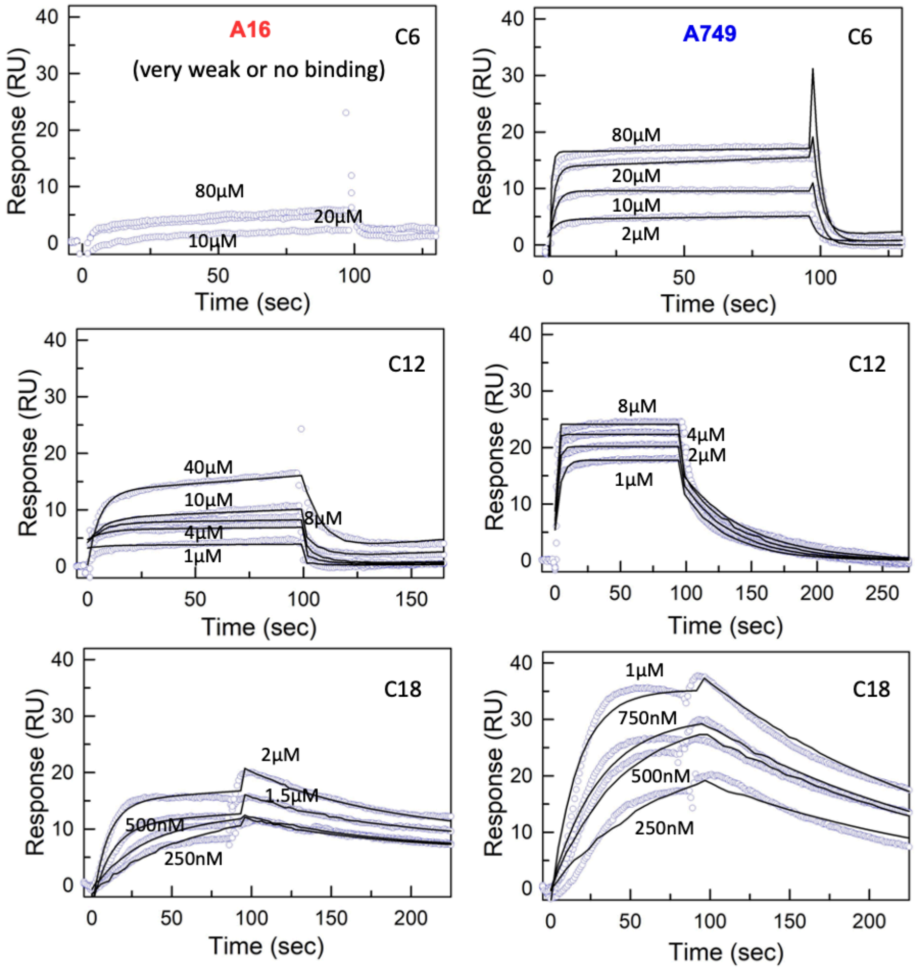


**Figure S12** SPR sensograms of different ligands binding to and unbinding from A16 (left column) and A749 (right column) at pH 7.4. Note the extremely slow unbinding of C18 from both the paralogs. The middle row is also shown in main text panels Figure 8D and 8E. Black lines passing through the data represent global fit from a 1:1 binding model.

**Supporting References**


(1) Crooks, G. E.; Hon, G.; Chandonia, J.-M.; Brenner, S. E. WebLogo: A Sequence Logo Generator. *Genome Res.* **2004**, *14* (6), 1188–1190. https://doi.org/10.1101/gr.849004.

(2) Guzman-Casado, M.; Parody-Morreale, A.; Robic, S.; Marqusee, S.; Sanchez-Ruiz, J. M. Energetic Evidence for Formation of a PH-Dependent Hydrophobic Cluster in the Denatured State of Thermus Thermophilus Ribonuclease H. *J. Mol. Biol.* **2003**, *329* (4), 731–743.

(3) Gopi, S.; Aranganathan, A.; Naganathan, A. N. Thermodynamics and Folding Landscapes of Large Proteins from a Statistical Mechanical Model. *Curr. Res. Struct. Biol.* **2019**, *1*, 6–12.

(4) Naganathan, A. N.; Dani, R.; Gopi, S.; Aranganathan, A.; Narayan, A. Folding Intermediates, Heterogeneous Native Ensembles and Protein Function. *J. Mol. Biol.* **2021**, *433* (24), 167325. https://doi.org/10.1016/j.jmb.2021.167325.
